# Weakened prefrontal activation dynamics associated with slowed information processing speed in multiple sclerosis

**DOI:** 10.1101/2025.04.08.647709

**Authors:** Olivier Burta, Fahimeh Akbarian, Chiara Rossi, Diego Vidaurre, Marie Bie D’hooghe, Miguel D’Haeseleer, Guy Nagels, Jeroen Van Schependom

**Author notes:** Contributions O.B. conducted the analysis and wrote the manuscript. J.V.S. was the main supervisor of the work and helped write and review the paper manuscript. F.A., C.R., D.V., and G.N. provided support regarding the analysis methods and provided comments during the manuscript writing. M.B.D. and M.H. provided comments on the work. All authors approved the submitted version.

## Abstract

Information processing speed (IPS) is a core cognitive deficit in people with multiple sclerosis (PwMS). Previous efforts have associated IPS performance to frontal regions, but were constrained by limited temporal resolution. In this work, we employed a data-driven method, the time delay embedded-hidden Markov model (TDE-HMM), to identify task-specific states that are spectrally defined with distinct temporal and spatial profiles. We used magnetoencephalographic (MEG) data recorded while healthy controls and PwMS performed a cognitive task designed to capture IPS, the Symbol Digit Modalities Test (SDMT). The TDE-HMM identified five task-relevant states, supporting a tri-factor contribution to IPS: sensory speed (occipital visual detection and processing), cognitive speed (prefrontal executive and frontoparietal attention shift), and motor speed (sensorimotor). We observed reduced prefrontal and increased frontoparietal activation in PwMS, which correlated with offline SDMT performance. This work can drive future research for MS treatments targeting IPS improvements.

## 1. Introduction

Multiple sclerosis (MS) is a leading neuroinflammatory disease in young adults, affecting the central nervous system (CNS).^1^ MS lesions can occur in multiple locations of the CNS, causing a large variety of symptoms ranging from physical to cognitive impairments.^1,2^ About 50% of people with multiple sclerosis (PwMS) experience cognitive impairment^3^, affecting domains such as attention, long-term memory, and working memory.^4^ However, information processing speed (IPS) stands out as a core cognitive deficit in PwMS.^5^ IPS is the cognitive domain most widely affected by MS^6^ and is one of the first deficits to emerge.^7^

The standard cognitive test used to capture IPS deficits is the Symbol Digit Modalities Test (SDMT).^8,9^ While the original paper-based test remains the standard in clinical practice, computerised adaptations are increasingly being investigated.^10^ Over the last decade, researchers have turned to using the computerised SDMT as a task paradigm in functional neuroimaging studies. When studied with functional magnetic resonance imaging (fMRI), PwMS showed larger recruitment and more connections to frontal areas during a computerised SDMT.^11^ Another fMRI study in people with relapsing-remitting MS (RRMS) found the frontal, superior parietal, occipital, and medial posterior cerebellar areas to activate.^12^ Compared to healthy controls, Grothe et al. also reported a decreased activation of the cingulate cortex in PwMS during execution of the SDMT.^12^ However, fMRI is not the optimal functional neuroimaging modality for studying the cognitive processes required during the SDMT due to its temporal unspecificity and being an indirect measure. With a temporal resolution in the range of seconds, fMRI is unable to capture the highly dynamic brain activity involved. As a consequence, the exact neurophysiological processes underlying SDMT execution, particularly how fast they unfold over time, remain unclear.

Electrophysiological brain activity can be measured at a much higher temporal resolution. Electroencephalography (EEG) and magnetoencephalography (MEG) capture the electric and magnetic fields generated in the brain as a result of neural activity, respectively. MEG studies revealed associations between partial functional disconnection of the temporal brain regions and overall cognitive capacity in MS.^13^ However, no study has characterised the brain networks underlying a task primarily targeting IPS while benefiting from the temporal resolution of M/EEG.

A powerful approach to explore and characterise transient fast-switching brain dynamics is based on Hidden Markov Models (HMMs).^14,15^ The HMM describes the time series using a set of states and the transitions between them. Specifically, using the time delay embedded-hidden Markov model (TDE-HMM), those hidden states represent transient and spectrally defined fast-switching functional brain networks.^16^ While traditional functional connectivity (FC) analysis methods often fall short in capturing the combined temporal, spectral, and spatial information^17^, the TDE-HMM successfully captures this multi-dimensional complexity. This state-of-the-art method has previously been applied in both resting-state^14,16,18^ and various task paradigms.^15,19–21^ In rest, Vidaurre et al. extracted a frontal and posterior default mode network (DMN) from MEG data.^16^ Using task data enables us to identify the functional brain networks underlying domain-specific cognitive processes. In this context, Rossi et al. unravelled the brain dynamics underlying working memory (WM) in a visual-verbal n-back task^19^, showing consistency with existing theoretical models of WM. By further including PwMS in the analysis, they were able to identify pathology-specific and treatment-induced effects captured in the brain states’ properties.^22^

In this work, we use the TDE-HMM method to study MEG brain activity during a task primarily assessing IPS, the SDMT. Our goal is to provide the first description of the neurophysiological processes underlying an IPS task. Further, by including an MS cohort, we aim to identify altered brain dynamics in PwMS, hypothesising that these will be disrupted and likely slowed compared to healthy controls. Finally, we demonstrate how our findings can lend support to existing theories of IPS deficits in MS.

## 2. Methods

### 2.1 Participants

The dataset consists of 110 subjects, 37 healthy controls (HCs) and 73 people with multiple sclerosis (PwMS). The MS cohort, diagnosed according to the 2010 McDonald criteria^23^, was recruited at the National MS Center Melsbroek, Belgium. It comprises 63 patients with relapsing-remitting MS (RRMS), 6 with primary progressive MS (PPMS), 3 with secondary progressive MS (SPMS), and 1 with clinically isolated syndrome (CIS). Exclusion criteria included an Expanded Disability Status Scale (EDSS) score greater than 6, or the presence of other neuropsychiatric disorders. Some subjects in the MS cohort are prescribed benzodiazepine (BZD) treatment, known to alter functional brain dynamics.^24^ This dataset contains 18 PwMS with (BZD+) and 55 PwMS without BZD treatment (BZD-). Ethical approval for data collection was provided by the ethical committees of the National MS Center Melsbroek (2015-02-12) and the University Hospital Brussels (Commissie Medische Ethiek UZ Brussel, B.U.N. 143201423263, 2015/11), while retrospective data analysis received ethical approval from the ethical committee of the University Hospital Brussels (B.U.N. 1432025000017, 2025/02).

All participants provided written informed consent and completed a paper version of the SDMT as part of the Brief International Cognitive Assessment in MS (BICAMS)^25^ on the MEG session day.

### 2.2 MEG and MRI data

At the start of MEG data collection, a 306-channel Elekta Neuromag Vectorview system in the Laboratoire de Cartographie Fonctionnelle du Cerveau in Erasme Hospital (Belgium) was used (33 subjects). Afterwards, the MEG scanner was upgraded to an Elekta Neuromag Triux scanner (77 subjects). Both scanners sampled the MEG signal at 1000 Hz within a 0.1-330 Hz frequency band. The 306 sensors are arranged in 102 triplets (1 triplet = 1 magnetometer, 1 planar and 1 axial gradiometer). During subject preparation, the Head Position Indicator (HPI) coil locations, the position of three fiducial points and the head shape were saved using a three-dimensional digitiser (Polhemus). MEG was recorded in a magnetically shielded room (Elekta, MaxShield) to reduce external magnetic interference. Dedicated electro-oculography (EOG) and electrocardiography (ECG) channels were used to measure ocular and cardiac-related activity.

Structural brain information was collected using magnetic resonance imaging (MRI). Scans were acquired using a 3T Philips MR system at 1x1x1 mm³ resolution with T1-weighted sequence using the following parameters: TR = 4.93 ms, FA 8°, 230 x 230 mm² FOW, 310 sagittal slices, leading to a 0.53 x 0.53 x 0.53 mm³ resolution. MRI acquisition was performed last to avoid MRI interference with the MEG recording, with a median delay of 5 days between both acquisitions.

### 2.3 SDMT paradigm

During MEG acquisition, subjects performed a computerised SDMT. The key is displayed on a screen, showing the correct symbol-digit matches. When the task begins, a first symbol-digit pair is shown, representing a single trial. The subject has at most 6 seconds to submit a response by pressing one of two buttons (correct/incorrect), indicating whether the shown pair matches one from the key. Subjects were asked to complete the task as fast as possible while minimising mistakes. This task design follows the same outline as proposed by Genova et al.^26^, but the paradigm used in the present study did not have a fixed interstimulus interval of 3 s. Figure 1 illustrates the task.

**Figure 1.**
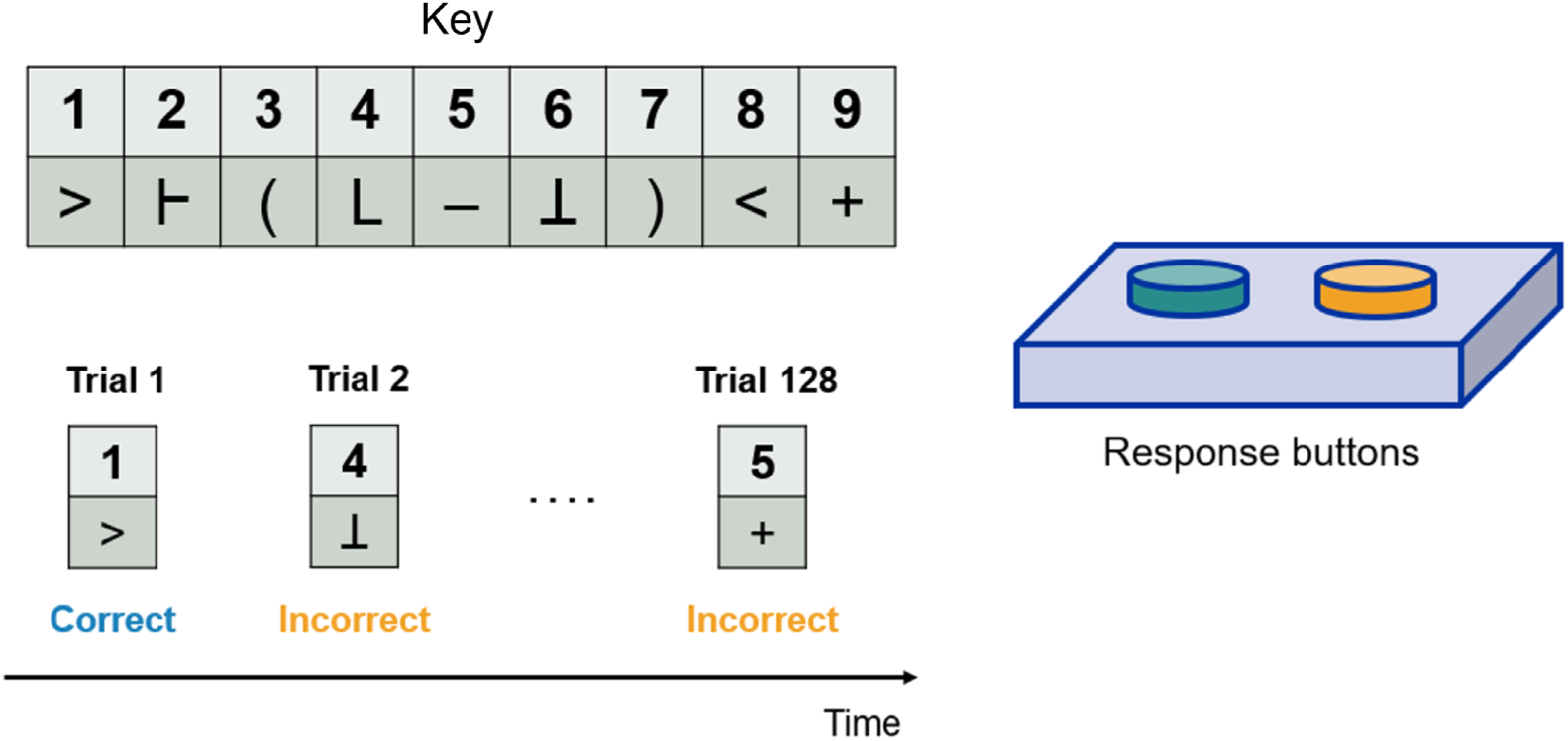
Graphical representation of the workflow during the computerised SDMT. Visual stimuli are presented as symbol-digit pairs. The subject is asked to press the blue button when the presented stimulus matches with the key (correct paradigm condition), and the orange button in the opposite case (incorrect paradigm condition).

Task performance was assessed using two behavioural metrics: reaction time (RT) and success rate. The RT is the exact time delay between stimulus presentation and the first recorded button press. The timing of visual stimulus onsets was saved using a photodiode setup located at the bottom of the projector screen. Success rate is calculated as the number of correct responses divided by the total number of trials.

Each participant first took a short training to ensure they understood the paradigm. The actual experiment consisted of 128 stimuli, split between 61 correct and 67 incorrect symbol-digit matches. Subjects did not receive feedback on their performance during or after the task.

### 2.4 MEG data processing

Before storing the recorded continuous MEG data, the MaxFilter software (Elekta, MaxFilter, version 2.2) transformed the data by compensating for close-by-artefacts and head movement using a temporal extension of the Signal Space Separation method (tSSS).^27^

#### Data preprocessing

All data processing steps were performed using the Oxford Centre for Human Brain Activity (OHBA) Software Library (OSL) package written in Python by the OHBA Analysis Group, University of Oxford, UK.^28^ The pipeline was applied to each subject separately. First, the data was downsampled to 250 Hz and filtered using a 5th-order Butterworth IIR bandpass filter from 0.01-70 Hz. This filtering operation removes non-neurophysiological signals, but we also applied an additional notch filter centred at 50 Hz (width of 2 Hz) to suppress artefactual power line interference. Next, we identified both data segments and recording channels of bad signal quality through a generalised extreme Studentised deviate (ESD) test at a significance level of 0.1. Each retrieved segment of bad quality (forming a window of 500 samples) was annotated in the data. The proportion of automatically determined bad channels is reported in the supplementary materials (section 1, Table S1). To remove remaining artefactual components, an automated independent component analysis (ICA) with dimension 64 was run. The independent components with the highest correlation to EOG/ECG channels were removed. Afterwards, the channels previously labelled as bad were reconstructed through spherical spline interpolation.^29^

#### Source reconstruction, parcellation, and sign-flipping

Before source reconstruction, each subject’s Polhemus headshape coordinates (acquired during MEG) were co-registered to their T1 MR scan using RHINO. An iterative process computes the optimal affine transformation between two spaces. Next, we used the Linearly Constrained Minimum Variance (LCMV) Beamformer to transform the preprocessed MEG data from sensor to source space.^30^ The source reconstruction was carried out using a single-layer forward model on an 8 mm uniform dipole grid.

To reduce dimensionality in source space, the source-reconstructed data is parcellated into 38 cortical brain regions using an atlas derived from an ICA of fMRI data from the Human Connectome Project, previously used in ref^31^. Using the spatial basis method, the time series attributed to each parcel corresponds to the first principal component from all voxels’ time series within that parcel, weighted by the spatial map’s coefficients. To reduce the effect of magnetic field spread, we applied a symmetric multivariate leakage correction algorithm to the parcelled data.^31^ This orthogonalisation procedure removes correlations at zero phase lag, while preserving non zero lag interactions for functional connectivity analyses.

So far, all processing steps were applied to single-subject data. MEG source reconstruction is characterised by inherent dipole ambiguity, posing a problem for analyses where all subjects’ data is concatenated. To counter this, we applied the sign-flipping algorithm proposed by Vidaurre et al. (parameters: parcels to flip = 20, number of iterations = 500).^21^

### 2.5 The Time delay embedded - hidden Markov model (TDE-HMM)

The Hidden Markov Model (HMM) describes the probabilistic activations of, and transitions between, a set of discrete states that best describe the observed MEG data. Together with their temporal activations, the HMM inference estimates the spatio-spectral properties of the states, which here represent functional brain networks. The Markovian assumption means that, if we knew which state was active at time points *t*-1 and *t*+1, the state activation at time point *t* would be conditionally independent of all the other time points, given *t*-1 and *t*+1.

Proven to be successful in describing brain dynamic patterns in other task paradigms^15,19^, we used the TDE-HMM implementation from the OSL Dynamics Toolbox in Python^32^. In essence, the TDE-HMM employs a Gaussian distribution on a time-delay embedded (TDE) space. This consists of shifting the original data over a set number of lags (here, N=15, spanning lags -7 to +7), allowing the incorporation of past and future time points into each step. Next, the matrix dimensionality is reduced by applying PCA, with the number of principal components at least twice that of the number of parcels (e.g., 38 x 2 = 76).^16^ Afterwards, the data is standardised (i.e. normalised to zero mean and unit variance). With these steps completed, the different model parameters (hidden states time course, transition probability matrix, and observation model) can be learned. These parameters are computed through stochastic variational Bayesian inference^21^, with updates for small batches of training data to save computations. To ensure that the HMM does not infer parameters from unreliable time series data, the previously labelled bad quality segments were omitted.

The main output consists of the inferred hidden states’ time course, providing the probability of each state activating at a given time point in the original MEG data. Subsequently, the time course is one-hot encoded, where each time point is assigned to the state with the highest activation probability, providing the input for subsequent analyses.

When using the TDE-HMM, a pre-specified number of states must be chosen. There is no golden standard for this parameter, it is typically adjusted based on the executed task. In this work, we ran five inferences with 4, 5, 6, 8 and 10 states to assess the influence of this parameter on the provided description.

### 2.6 State-wise temporal description

We applied the TDE-HMM to investigate how the different states (de)activate during SDMT execution. We performed an event-related (ER) analysis on the hidden state time courses. The time course was epoched between [-500, 3000] ms relative to stimulus onset, given an average RT at around 1.75 s post-stimulus for this task. If an epoch contained a segment of bad signal quality (section 2.4), that epoch was rejected. The proportion of rejected epochs is reported in the supplementary materials (section 1, Table S1). Remaining trials were baseline-corrected using the pre-stimulus [-500, 0] ms data. Afterwards, trials of the same stimulus type (correct/incorrect) were averaged together. This was repeated for each state and subject separately.

To determine whether the inferred states are task-related, we tested the statistical significance of the ER activation profile using a two-level generalized linear model (GLM). This analysis was performed using the *evoked_response_max_stat_perm()*^32^ function from osl-dynamics, offering flexibility to incorporate covariates. GLM input included 220 entries, with one subject-averaged epoch (110 subjects) for each stimulus type (2 conditions). The GLM design matrix consisted of 5 regressors: a constant regressor (mean activity) and four categorical regressors (stimulus type, presence of disease, benzodiazepine use, and gender). At the second level, we fitted individual contrasts of parameter estimates (COPES) at the group level to assess overall task-related activation, as well as the influence of covariates. Significance of state activations in the Mean Effect contrast was evaluated using a maximum statistic permutation test (N=1000) across time, states, and regressors. The maximal t-statistic was used to correct for multiple comparisons by controlling the familywise error rate, a common procedure in functional neuroimaging studies.^33^ The significance level was set to 1%. Using the same statistical procedure, we evaluated four contrasts to examine how the categorical regressors modulate state activations, for instance testing at each time point whether HCs and PwMS exhibit significantly different activation profiles.

For states where the disease effect contrast turned out as significant, we conducted a peak analysis. First, we extracted the maximum peak amplitude and latency within given time windows from the subject-averaged epochs, and then tested for significant differences between HCs and PwMS using a permutation test (t-statistic, N=1000, False Discovery Rate (FDR) correction for multiple comparisons). In states with significant peak differences, we investigated correlations (Pearson’s correlation) between the peak features and behavioural data, considering all subjects. The behavioural data consisted of mean RTs and offline SDMT z-scores. Correlations were corrected for multiple comparisons (via FDR), with 8 correlations in total, calculated as 1 state activation peak x 2 peak features (amplitude and latency) x 2 behavioural measures x 2 paradigm conditions (correct/incorrect).

### 2.7 Spectral and spatial characteristics

The spectral properties of the states are extracted using a non-parametric multitaper spectral estimation applied to every subject and state separately.^34^ This method takes the original parcelled MEG data multiplied by the hidden state time course as input, and produces power spectral densities (PSD) as output. These PSDs were originally determined for a wideband frequency range of [1-45] Hz. Subsequently, we estimated the state-specific spatial description by integrating the wideband PSDs over four frequency bands extracted using non-negative matrix factorisation (NNMF).^15^ These data-driven frequency bands are referred to as spectral components. To decide which spectral component best describes the spectral properties of a certain state, first, the mean PSD and mean coherence (averaged across all subjects and brain parcels) were plotted as a function of frequency, for that state. Next, similarity was sought in the frequency ranges between each data-driven spectral component and the frequency range of the strongest peak in both mean PSD and mean coherence plots. To plot the PSDs as spatial maps, each state-specific PSD was weighted with the most relevant spectral component for that state. We then took the weighted power, averaged it across all subjects, and report the spatial profiles as a z-score PSD distribution across brain parcels.

To quantify spectral synchronisation between brain regions, the PSDs were used to compute the coherence at each frequency bin, as explained by Vidaurre et al.^16^ The obtained coherence networks are state-specific, scaled by the most relevant spectral component for each state. For visualisation, only the connections remaining after Gaussian mixture model (GMM) thresholding are shown on both a circular plot and a brain mask.

## 3. Results

### 3.1 Behavioural data

The demographic and clinical characteristics of different subgroups in this study are shown in Table 1. The HC and PwMS groups are matched on gender (*χ*^2^ test, p-value = 0.18) and age (Mann-Whitney U-test, p-value = 0.84). The mean education level (Mann-Whitney U-test, p-value = 0.005) and SDMT score (Mann-Whitney U-test, p-value = 0.007) are significantly higher for HCs compared to PwMS in this dataset. The BZD+ and BZD-groups within the MS cohort significantly differ in terms of gender (Fisher exact test, p-value = 0.004) and EDSS score (Mann-Whitney U-test, p-value = 0.040). The success rates and RTs during the computerised SDMT are reported in the supplementary materials (section 2, Figure S1 and Table S2).

**Table 1.**
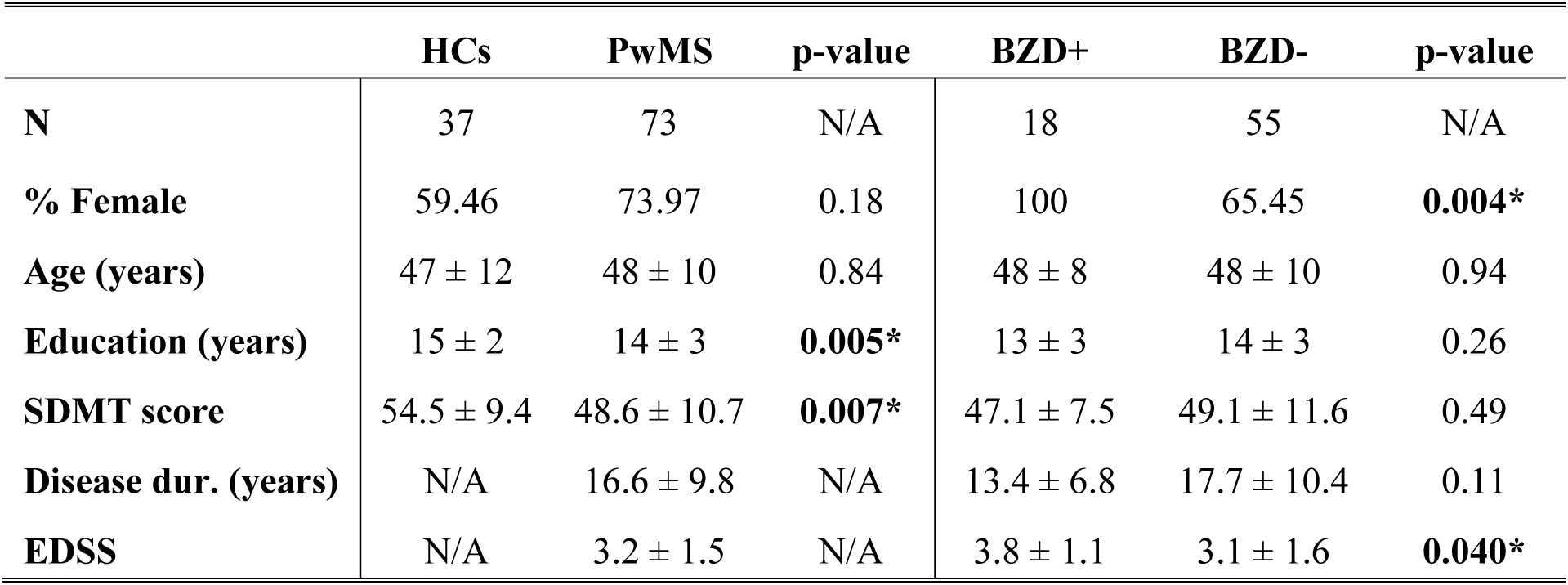
Demographic and clinical characteristics of the SDMT dataset. Statistical comparisons were conducted between healthy controls (HCs) and people with multiple sclerosis (PwMS), with an additional group comparison between BZD+ (PwMS with BZD treatment) and BZD- (PwMS without BZD treatment). Age, education, SDMT score (on-paper Symbol Digit Modalities Test), disease duration, and EDSS (Expanded Disability Status Scale) are reported as mean ± std. Differences in continuous outcomes across groups were compared using the Mann-Whitney U-test. N/A = not applicable, *p < 0.05.

### 3.2 Model inference

The TDE-HMM was run five times to infer 4, 5, 6, 8 and 10 states from the MEG data separately, allowing to assess the model’s robustness.^35^ The choice of hyperparameters for both the data preparation (time embedding, PCA) and the inference process can be found in the supplementary materials (section 3, Table S3). We observed that the 8 and 10 states inference provided some states with overlapping temporal and spatial characteristics. While further increasing the number of states may lead to some highly similar states, reducing the number of states may offer a too coarse solution, with distinct functional networks collapsing into the same state. In our data, we found that the 6-state solution strikes a good balance, and the results from this solution are shown and discussed.

The stochastic nature of the variational Bayesian inference process^21^ requires us to assess the stability and replicability of our results. Therefore, we trained the 6-state model four separate times. We determined the model inference to be stable, as states with similar spatiotemporal characteristics were consistently retrieved. Additionally, several summary statistics, describing the general dynamics of the states’ activation over the entire inference, are reported in the supplementary materials (section 4, Figure S2 and Table S4).

### 3.3 Task-related brain states

Figure 2 shows the average task-evoked state activation profiles during the computerised SDMT. The task-relevant states are those that are significantly activated or deactivated throughout the considered epoch window: states 1, 2, 3, 5, and 6 appear to be task-related. State 4 reflects lateral occipital baseline activity with properties (summary metrics, spatial profile) consistent with TDE-HMM literature^15^, and its description is reported in the supplementary materials (section 7, Figure S8). Remaining GLM contrasts (stimulus type effect, disease effect, benzodiazepine effect, and gender effect) are reported in the supplementary materials (section 5, Figure S3).

**Figure 2.**
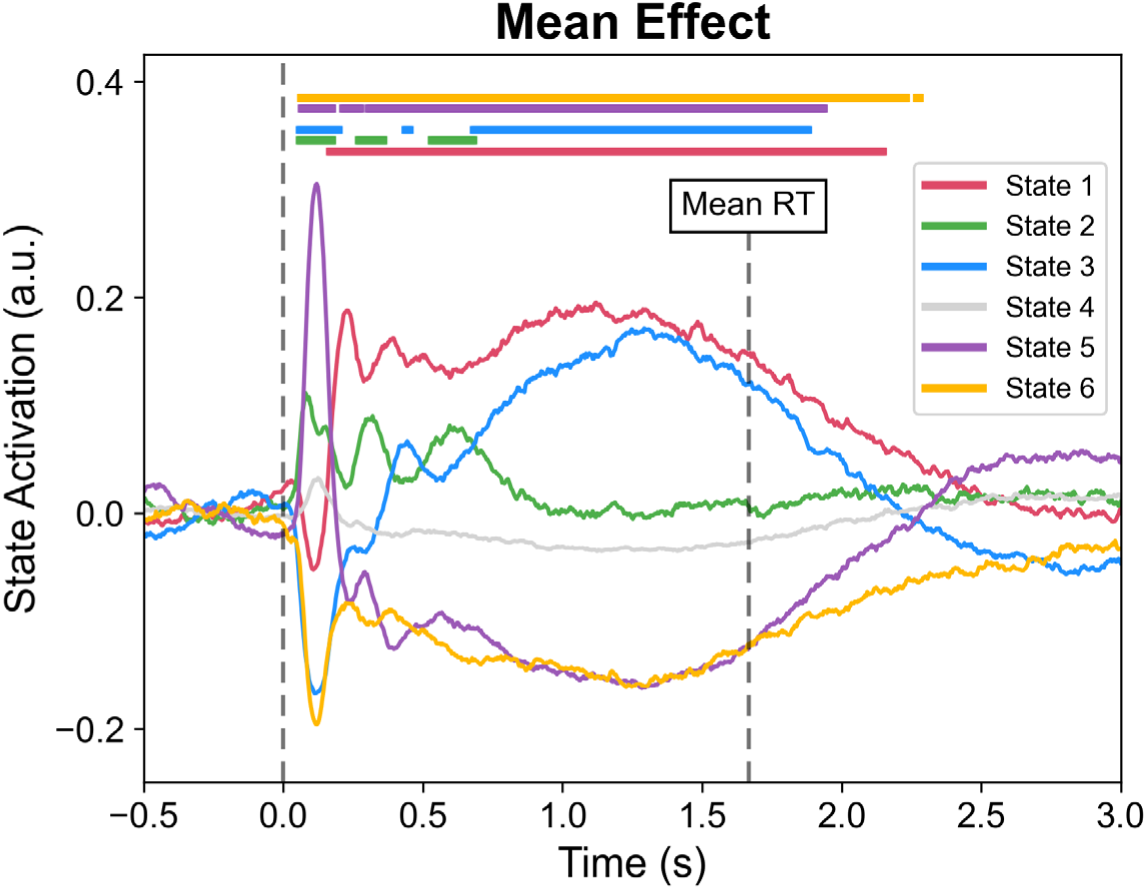
Average event-related activation for all states. Each curve – associated to one state – represents the average brain response, considered within [-500, 3000] ms around the stimulus onset, across subjects and paradigm conditions (provided by the mean effect contrast, GLM statistics). The horizontal lines at the top indicate time points of significant state activation or deactivation, determined using a permutation test (N = 1000) with the maximum statistic (corrected for multiple comparisons across states and time points) at a significance level of 1%.

### 3.4 Spectral contents

Four spectral components were extracted from the data, with their profiles reported in the supplementary materials (section 6, Figure S4). These findings are consistent with those reported in previous TDE-HMM studies.^16,19^ The three relevant spectral components, primarily representing the low-frequencies (1-8 Hz), alpha band (8-13 Hz), and beta band (13-30 Hz), were used to extract the weighted state-specific spatial maps and spectral coherence networks.

The state-specific mean PSD (Figure 3a) and mean coherence (Figure 3b) across all subjects were used to determine which spectral component best describes the properties of a particular state.

**Figure 3.**
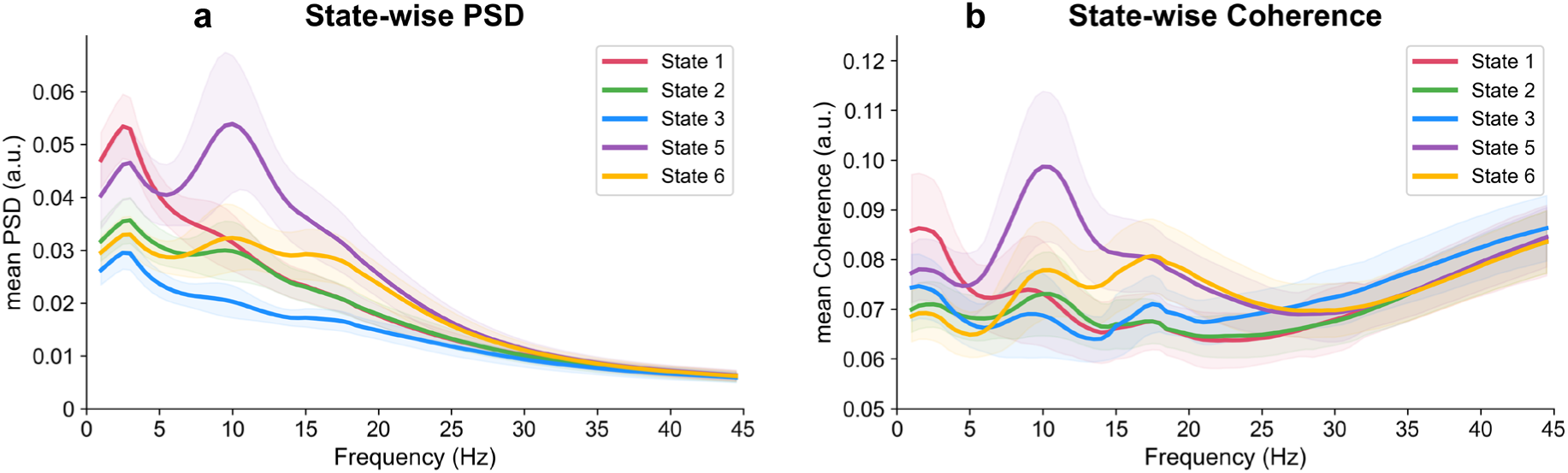
State-wise (a) mean power spectral density (PSD) and (b) mean coherence for only the task-relevant states. PSD and coherence are averaged across subjects, and across all regions (PSD) or pairs of regions (coherence). Shaded areas at each frequency bin correspond to the standard deviation across subjects.

### 3.5 Description of brain states

#### State 1

State 1 shows a task-related response significantly activated between 250 and 1250 ms post-stimulus (Figure 2). After that, the mean activation probability gradually decreases until 2250 ms post-stimulus. The activity of state 1 is mostly contained within spectral component 1, as can be seen from the low-frequency peak in both PSD (Figure 3a) and coherence (Figure 3b). From the z-scored PSD map, mainly frontal regions show the highest power. The coherence networks reveal the involvement of prefrontal (e.g. superior, ventrolateral, orbitofrontal), medial sensorimotor, posterior cingulate cortex (PCC), and anterior temporal regions.

#### State 2

State 2 significantly activates at 300 ms post-stimulus, with a second significant peak around 550 ms post-stimulus (Figure 2). Distinct alpha peaks in both mean PSD (Figure 3a) and coherence plots (Figure 3b) lead to this state being associated with spectral component 2. The spatial PSD map highlights the occipital lobe. This is further supported by the coherence networks showing high synchronisation between several regions in the occipital brain area. More specifically, this concerns the inferior occipital, middle occipital, occipital pole, and cuneus regions, with multiple connections spanning across the two hemispheres. For TDE-HMM runs with a lower number of hidden states (k=4, k=5), state 2 and state 5 collapse into a single occipital state.

#### State 3

State 3 is characterised by a significant increase in activation at 450 ms post-stimulus, followed by sustained activation between 750 and 1500 ms (Figure 2). The significant state deactivation around 100 ms post-stimulus is likely an artifact of the mutual exclusivity assumption (i.e. other states, state 2, and 5 in the presented results, significantly activate, causing the remaining states to deactivate w.r.t. baseline). Therefore, this specific peak is unlikely to be task-related to state 3. On the spatio-spectral level, state 3 represents a frontoparietal network characterised by broadband coherence. While the mean PSD (Figure 3a) appears rather flat across all frequencies, the mean coherence (Figure 3b) displays small peaks for each spectral component’s frequency range. With the beta range peak standing out the most, we determined state 3’s spectral content to be mainly focused on beta activity. State 3 activates in the frontal (ventrolateral and dorsal prefrontal cortex) and parietal (supramarginal gyrus) lobes. Coherence networks display broad connectivity patterns, with consistently retrieved parieto-occipital coherence in all three spectral components (Figure S7).

#### State 5

State 5 is characterised by the strongest significant activation between 50 and 100 ms post-stimulus across all states (Figure 2). This is followed by significant deactivation between 300 and 1750 ms post-stimulus. Similarly to state 2, state 5 is characterised by alpha activity (spectral component 2) in the occipital lobe. Its coherence network is similar to that of state 2, although the connectivity strength in state 5 is overall stronger.

#### State 6

State 6 significantly deactivates between 100 and 1300 ms post-stimulus, after which reactivation builds up (Figure 2). This reactivation occurs about 450 ms before the mean RT to the stimuli (1750 ms post-stimulus). Based on a peak in the PSD (Figure 3a) and the strongest coherence peak (Figure 3b), both in the beta frequency range, the main spectral content of state 6 is summarised by spectral component 3. The spatial PSD map of state 6 displays activity in the supramarginal sensorimotor cortex, present in both hemispheres. On top of the regions identified through the spatial map, the coherence networks display strong connections between the left and right superior prefrontal, and the left and right angular gyri. Notably, the lateral and medial sensorimotor cortices show inter-hemisphere connections.

### 3.6 Group-level differences and correlations

#### Temporal state activation differences

Figures 4 and 5 highlight time points where the HCs and PwMS temporal activation profiles significantly differ (max t-stat permutation, N = 1000, p < 0.05). State 1, representing a prefrontal network, has a significantly increased activation in HCs around 200-250 ms post-stimulus (Figure 4a). The frontoparietal network (state 3) exhibits a significantly increased activation in PwMS across most of the epoch window (Figure 4e). Lastly, the occipital state 5 shows significantly reduced activation in PwMS between 50-100 ms, 450-650 ms, and 1300-2200 ms post-stimulus (Figure 5a). Modulation of the states’ activation profiles by other covariates, including benzodiazepine treatment and gender, is discussed in the supplementary materials (section 5).

**Figure 4.**
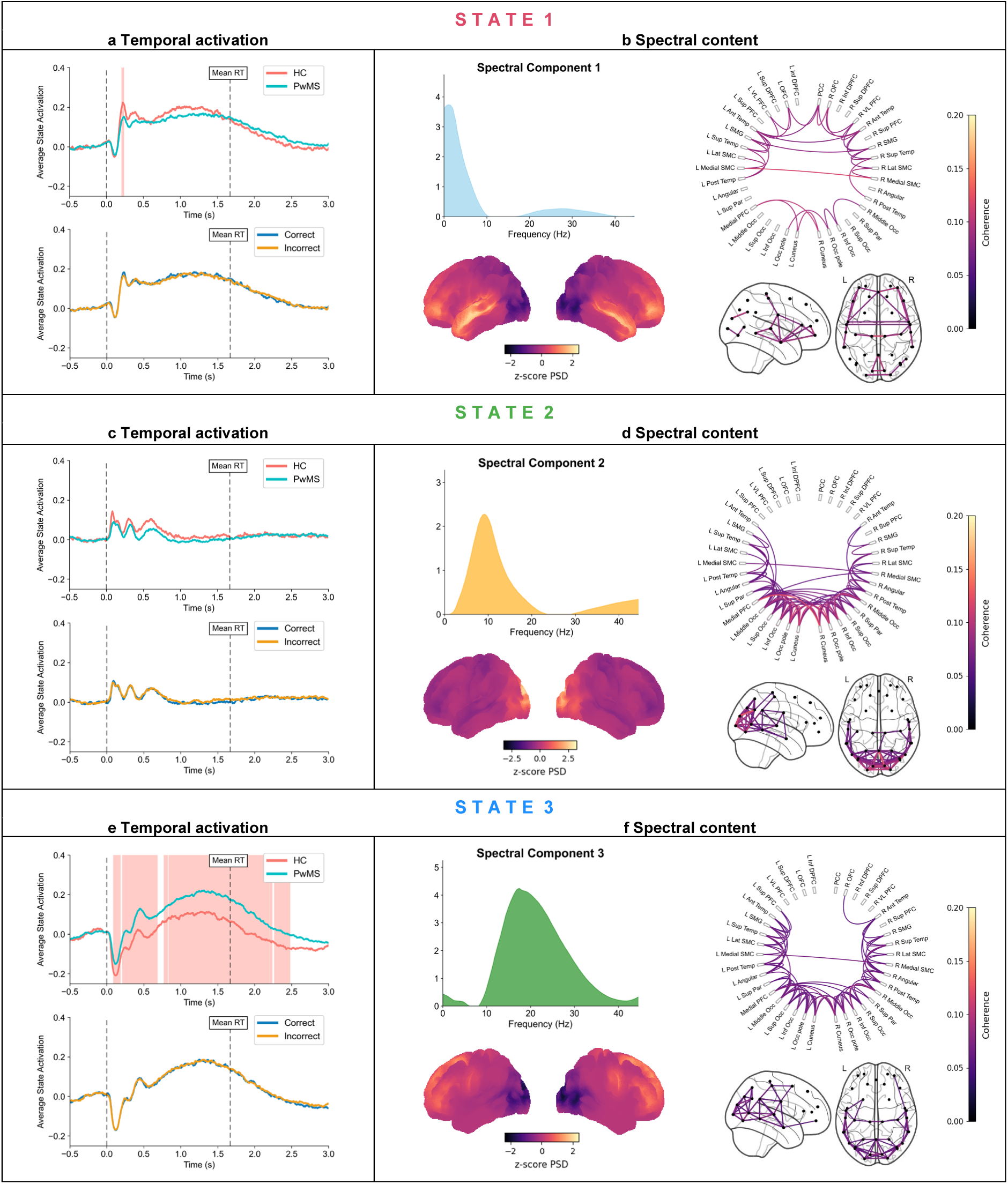
Description of state 1, 2, and 3. We report the (a, c, e) temporal activation pattern and (b, d, f) spectral content of each state. The temporal activation profile plots for (a) state 1, (c) state 2, and (e) state 3 each display two comparisons: disease group (HCs vs. PwMS) and paradigm condition (correct vs. incorrect), with solid lines representing the group-averaged epochs. Dotted vertical lines indicate the stimulus onset and mean reaction time, while pink shaded areas highlight time points where GLM contrasts reveal significant differences across groups or task conditions (max t-stat permutation, N = 1000, p < 0.05). Next, the spectral content boxes of (b) state 1, (d) state 2, and (f) state 3 contain: the spectral component best associated with the state, the group-level average PSD map, and the coherence network. The z-score PSD (standardised across parcels) was computed within the relevant spectral component’s weighted frequency range. Lastly, the coherence networks are shown as circular plots and through brain masks. Only connections surviving thresholding via GMM are displayed.

**Figure 5.**
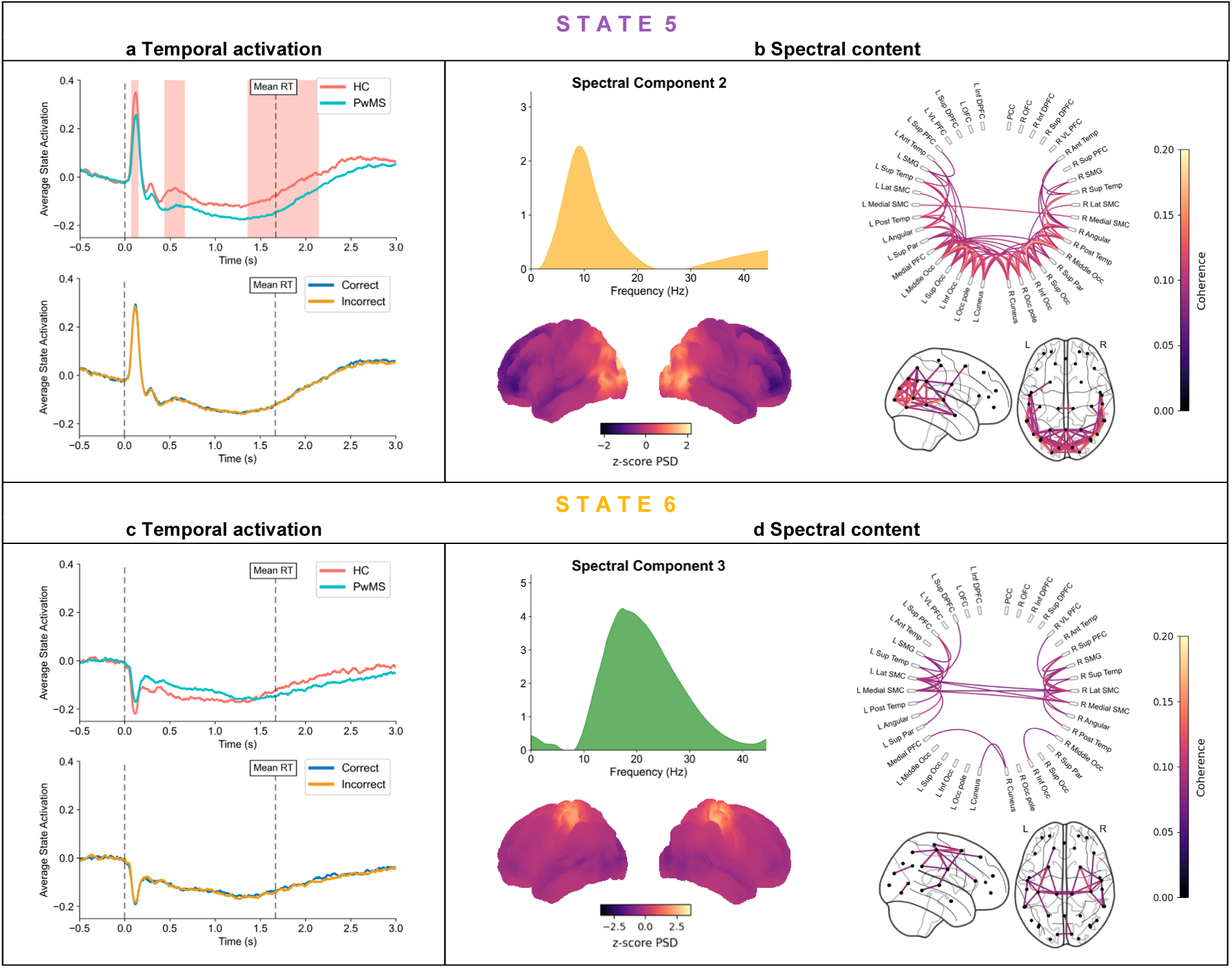
Description of state 5 and 6. We report the (a, c) temporal activation pattern and (b, d) spectral content of each state. The temporal activation profile plots for (a) state 5 and (c) state 6 each display two comparisons: disease group (HCs vs. PwMS) and paradigm condition (correct vs. incorrect), with solid lines representing the group-averaged epochs. Dotted vertical lines indicate the stimulus onset and mean reaction time, while pink shaded areas highlight time points where GLM contrasts reveal significant differences across groups or task conditions (max t-stat permutation, N = 1000, p < 0.05). Next, the spectral content boxes of (b) state 5 and (d) state 6 contain: the spectral component best associated with the state, the group-level average PSD map, and the coherence network. The z-score PSD (standardised across parcels) was computed within the relevant spectral component’s weighted frequency range. Lastly, the coherence networks are shown as circular plots and through brain masks. Only connections surviving thresholding via GMM are displayed.

#### Peak analysis and correlations

Building on these results, we performed a peak analysis to compare the maximum amplitude and latency of early activation peaks in the states with significantly altered activation between HCs and PwMS (state 1, 3, and 5). Figure 6 shows the distributions of peak features extracted from the time windows 150-300 ms post-stimulus (state 1), 450-800 ms post-stimulus (state 3), and 50-200 ms post-stimulus (state 5). This analysis revealed a significantly reduced maximum amplitude of the early activation peak in state 1 for PwMS compared to HCs, across both paradigm conditions (Figure 6a). By correlating these peak features to behavioural measures from all subjects’ data, we found the max peak amplitude to significantly negatively correlate with the RT in both paradigm conditions (Figure 7). Additionally, the max peak latency of state 1 shows a non-significant positive correlation with the RT. Taken together, faster and more likely early prefrontal state activation is associated with shorter RTs during the SDMT paradigm. Moreover, the max peak amplitude in state 1 significantly positively correlates with the z-score SDMT, for both task conditions. Although the correlation analysis included data from all 110 subjects, similar findings emerged when considering the HC and PwMS subgroups separately.

**Figure 6.**
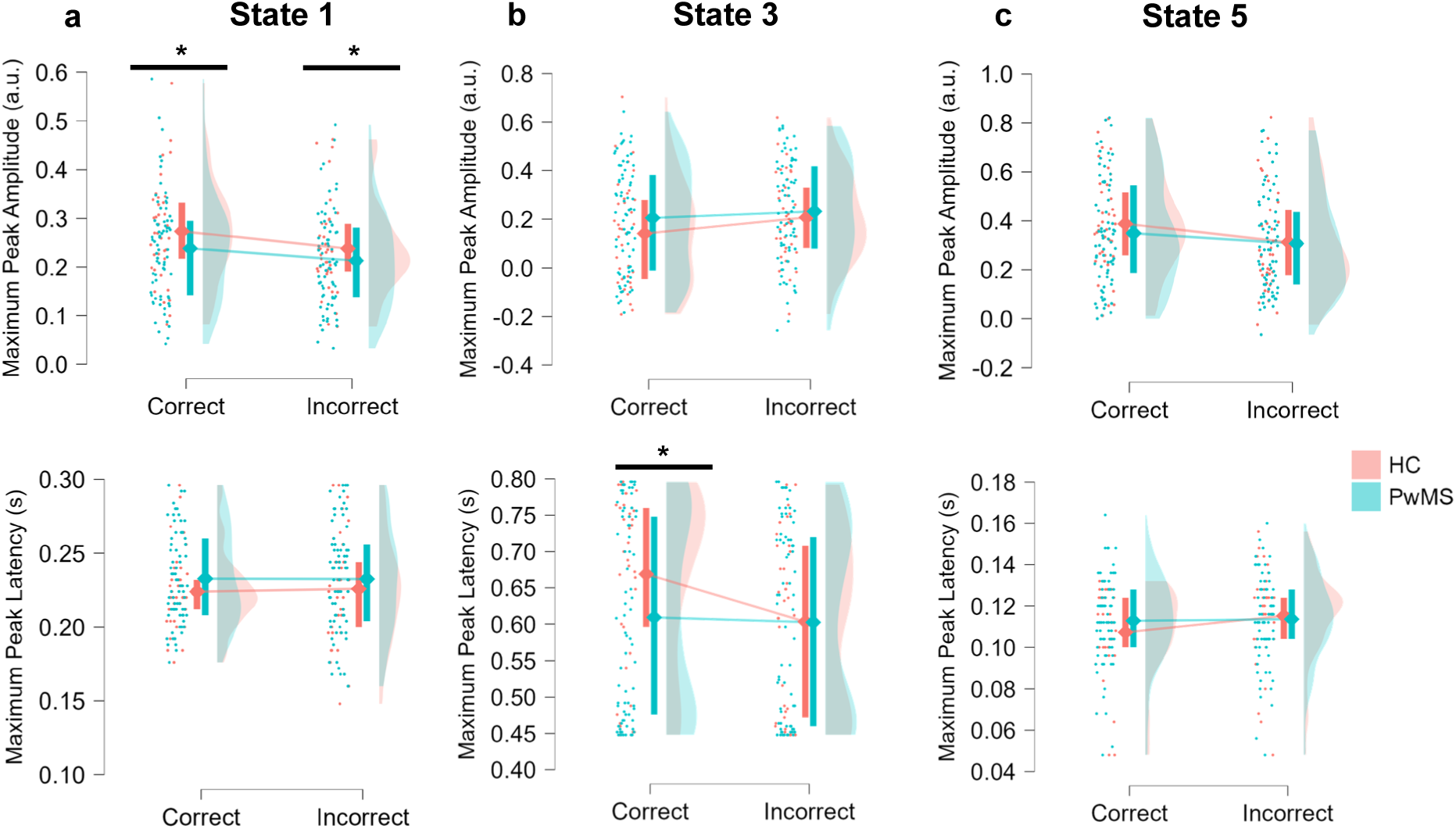
Distributions of event-related peak features for HCs and PwMS. Each subplot shows the distributions of peak amplitude or latency for features extracted at the maximum peak of (a) state 1, (b) state 3, and (c) state 5 in both groups (HCs and PwMS), with separate distributions by paradigm condition (correct/incorrect). Black horizontal lines indicate significant group differences (permutation t-test, FDR correction for 12 comparisons, *p < 0.05).

**Figure 7.**
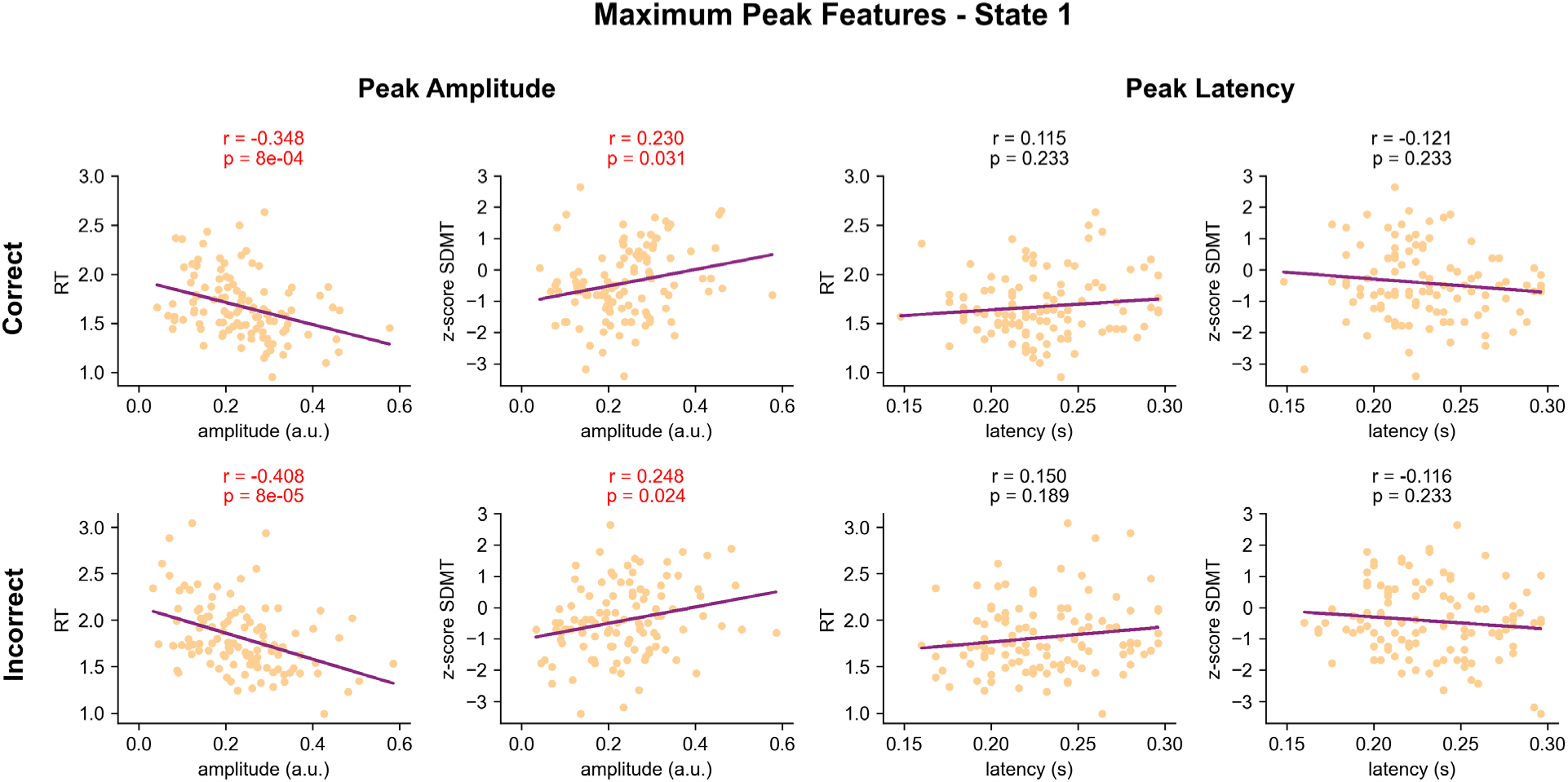
Correlations between state activations’ peak features and behavioural data. Peak features consist of amplitude and latency in the maximum peak of state 1. The behavioural measures include the mean reaction time (RT) per subject, and the standardised (z-score) SDMT. The relationships are shown as a scatter plot and regression line, whose correlation coefficient (r) is computed using Pearson’s correlation. The p-values (p) are corrected for multiple comparisons via FDR. Each row uses data from one of the two paradigm conditions (correct or incorrect).

In the frontoparietal state 3, we observe a significantly increased peak latency in HCs (Figure 6b). However, given the bimodal-like distribution of the maximum peak latency in state 3, we chose not to include this state in the subsequent correlation analysis. Finally, the initial activation of state 5 does not reflect significantly different peak measures between HCs and PwMS (Figure 6c).

## 4. Discussion

In this work, we used the TDE-HMM to extract data-driven brain states with distinct spectral profiles, involved during the execution of an SDMT task. So far, fMRI studies have consistently reported strong activation of the frontal, parietal, and occipital lobes in healthy individuals.^36,37^ In clinical populations with impaired IPS, such as PwMS, increased frontal recruitment and enhanced connectivity are particularly evident during early disease stages.^11^ Another study reported reduced cingulate cortex activation, interpreted as diminished attentional drive, in PwMS compared to healthy controls.^12^ However, these studies could not capture the fast temporal dynamics underlying SDMT execution. In the following, we discuss the extracted TDE-HMM states in their order of activation throughout the task.

Visual stimuli presented during the SDMT are first captured in the occipital cortex. State 5’s activation at a latency of 100 ms resembles the P100 visual-evoked potentials (VEPs) reported in literature.^38,39^ This occipital state is mainly characterised by alpha activity, previously associated with the initial detection of visual stimuli.^40,41^ The VEP literature repeatedly reports prolonged latencies and reduced amplitudes in the P100 responses of PwMS.^42,43^ Although we found a reduced temporal activation in PwMS compared to HCs (Figure 5a), our peak analysis did not reveal significant differences in peak amplitude or latency between these cohorts.

Following the initial stimulus detection, a prefrontal network activates at 250 ms post-stimulus and persists until 1500 ms post-stimulus. State 1 arises predominantly in the prefrontal cortex, consistent with fMRI findings.^12,36,37^ The prefrontal cortex plays a crucial role in numerous cognitive processes, including stimulus encoding, information integration, and goal-directed behaviour, as well as in emotional regulation.^44,45^ Most of this state’s spectral content is described within the low-frequencies, specifically the theta range. Given that prefrontal theta activity has been widely associated with memory encoding and higher-order cognitive processing^46,47^, this state could play a role in stimulus encoding during the SDMT. Another cognitive process often associated with executive frontal regions is that of decision-making.^48,49^ Considering that state 1 remains significantly activated until 300 ms before the mean RT, this could capture the response button selection near the end of an SDMT trial.

We found that PwMS show significantly reduced activation of this prefrontal network compared to HCs, only at early state activation (250 ms post-stimulus). Decreased prefrontal activation was previously reported using fMRI^26^, and more recently in a follow-up study by Rossi et al. using the same TDE-HMM approach.^22^ Impaired functional activation of state 1 could be a result of structural brain damage.^50^ As such, Grothe et al. have shown an association between IPS performance and structural white matter damage in frontoparietal regions in PwMS.^51^ Our work also revealed significant correlations between maximum peak features of prefrontal state activation and both task reaction times (online) and z-score SDMT performance (offline). This further suggests that the integrity of the executive control unit in the prefrontal cortex contributes to slowed IPS in PwMS.^52^

In the next phase of the SDMT, the presented stimulus is compared against the key. The extracted state 3, representing a frontoparietal network, may play a crucial role in this process. Previous studies have associated parietal activations with spatial attention shifts during visual tasks.^53,54^ Other research has linked functional connectivity between occipital and parietal regions with visual information processing.^55,56^ Given that our frontoparietal state is characterised by a parieto-occipital coherence network, this points to its involvement in visual stimulus processing during the SDMT.

This is reinforced by looking at the successive significant activations of state 2 (occipital) and state 3 (frontoparietal) depicted in Figure 2. Specifically, the occipital state first activates at 300 ms post-stimulus, followed by the frontoparietal state at 450 ms, and again occipital activation at 550 ms. In the context of the SDMT paradigm, occipital state activations likely reflect visual processing of the presented stimulus (300 ms) and later the key (550 ms), while the frontoparietal activation at 450 ms post-stimulus could support the spatial shift guiding the eye movement (saccade) between these targets. In addition, our results show that both states exhibit parieto-occipital coherence in the alpha band (Figures S6 and S7). This aligns with findings by Zhigalov et al., who suggest that alpha oscillations facilitate visual information gating between parieto-occipital regions^57^, supporting our interpretation.

Alongside alpha-coherence, this frontoparietal state is further marked by beta-coherence. Antzoulatos et al. propose that beta-band synchrony within frontoparietal networks is crucial for maintaining task-relevant representations, as demonstrated during a visual categorisation task.^58^ In the SDMT paradigm, frontoparietal activation between 750 and 1500 ms post-stimulus may reflect the mental processes involved in categorising the presented stimulus as either a correct or incorrect match to the key. This categorisation could then inform decision-making by the prefrontal network, as indicated by the simultaneous event-related activation patterns in both frontoparietal and prefrontal states. Interestingly, the main analysis (6-state model) revealed a frontoparietal network containing both alpha and beta coherence, which remained within a single state even when expanding the model to infer more states (k=10). This may suggest one of two potential explanations: that these processes are inherently interrelated, or that this reflects a methodological limitation of our approach.

Compared to HCs, PwMS exhibit a significantly increased temporal activation of the frontoparietal state, maintained across the full epoch window (Figure 4e). A review by Rocca et al. highlights increased frontoparietal fMRI activity in PwMS who maintain intact performance during cognitive tasks.^59^ This is consistent with the theory of compensatory network activations in neurological diseases^60^, where increased frontoparietal activity may reflect the brain’s attempt to sustain cognitive performance despite MS-induced dysfunction.

The final step in the task paradigm consists of reporting whether the presented stimulus matches the key by pressing one of two buttons. The spatio-spectral layout of state 6 corresponds to a typical sensorimotor network, characterised by beta activity in the left and right somatosensory cortices.^61^ Previous TDE-HMM studies reported similar states in MEG task paradigms requiring a motor response.^15,19^

Overall, we identified five brain states relevant to computerised SDMT execution, each having distinct temporal, spectral, and spatial characteristics. To relate this work to existing theoretical models of IPS, the tri-factor model of IPS in MS is used.^62^ Our analysis gives support to this model, disambiguating IPS as three contributing factors: sensory, cognitive, and motor speed. Combining the previously discussed results with the tri-factor model of IPS, a novel description of the neurophysiological processes underlying the SDMT is proposed in Figure 8.

**Figure 8.**
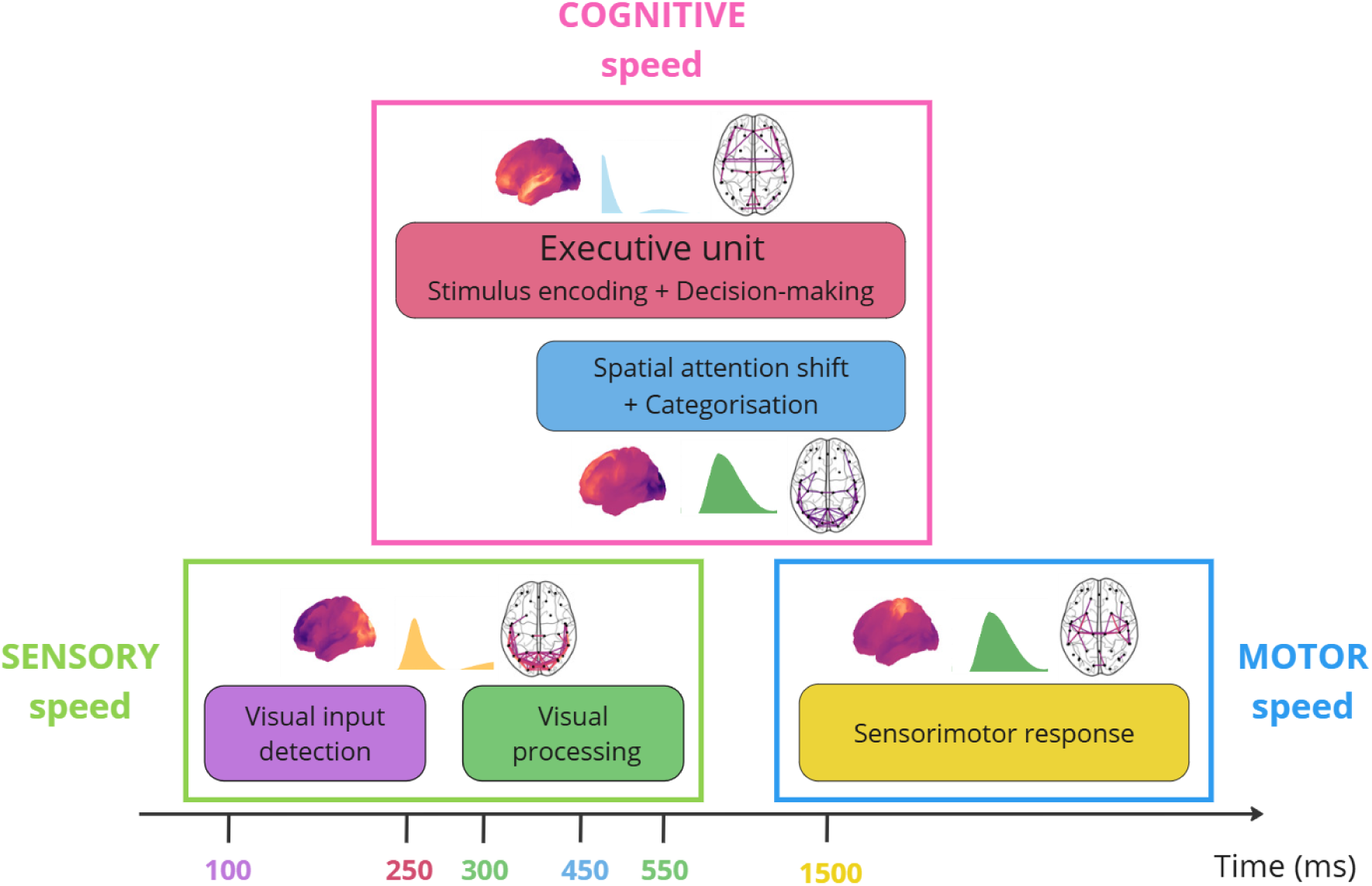
Summary of the proposed pathway of the neurophysiological processes unfolding during the computerised SDMT. The rectangles with filled colors and the numbers on the time axis are color-coded to match the respective states sharing the same colors in Figures 4 and 5. Using the tri-factor model of IPS in MS^62^, the pathway is divided into 3 parts: sensory, cognitive, and motor speed.

Although the SDMT is highly sensitive in capturing slowed IPS in MS, it should be acknowledged that the SDMT is not a pure measure of IPS.^63^ Aspects such as visual scanning or motor dysfunctions influence the assessment of IPS.^9^ This is particularly important when research focuses specifically on the cognitive impairments in neurological diseases such as MS. In theory, one could extract the pure cognitive speed component by isolating the sensory and motor components contributing to IPS assessed by the SDMT. On the one hand, the sensory speed can be measured through the latency of VEPs.^39^ On the other hand, the motor speed can be determined by means of transcranial magnetic stimulation (TMS) over the motor cortex.^64^ Future studies could explore this deeper by including additional experiments with the aim of isolating the targeted cognitive processing speed.

Previous work by Rossi et al. described the brain dynamics involved during a working memory (WM) task, the n-back.^19,22^ Comparing the state descriptions between the n-back (WM task) and the SDMT (IPS task) reveals several brain states shared between both cognitive tasks, as well as task-unique states. Both tasks share a prefrontal theta, sensorimotor beta, and occipital alpha states. Specific to the n-back, Rossi et al. identified an M300 state important for memory recall and response selection. In our work, we identified a frontoparietal network, crucial for supporting attentional shifts between visual targets, unique to the SDMT. Not only can different tasks capture different task-specific cognitive processes, but they can also highlight similar network dysfunctions in PwMS. As such, prefrontal executive unit activation was revealed to be impaired in PwMS during both the n-back and SDMT.

## 5. Limitations

In this work, we investigated MS-induced alterations in brain network activations within the extracted TDE-HMM states. Future studies should examine how these functional changes relate to structural MRI data. Given the key role of demyelination in IPS, incorporating MRI-derived structural measures could offer valuable insights into subject-specific disruptions of functional brain dynamics.

A limitation of the HMM is that the model assumes that only one state is active per time point, which may be too restrictive to fully represent brain function. Dynamic Network Modes (DyNeMo), proposed by Gohil et al.^65^, is a recently introduced technique that overcomes this limitation. Future studies could explore the DyNeMo approach to assess its potential to provide new insights regarding the network dynamics during execution of various cognitive tasks.

## Conclusion

This study provides a detailed description of the neurophysiological processes underlying an IPS task. We applied the TDE-HMM method to task-MEG data, recorded while HCs and PwMS performed the SDMT. This revealed five task-relevant brain states with distinct temporal, spectral, and spatial characteristics. By including MEG data from PwMS, we revealed a reduced activation of the early prefrontal theta executive unit and enhanced activation of the frontoparietal network in this population. This way, our findings lend support to the tri-factor model of IPS in MS, which proposes that sensory, cognitive, and motor speed components collectively influence IPS assessments. The observed impairments, alongside the novel detailed network descriptions, can drive future research for treatments targeting the improvement of IPS performance.

## Data Availability

The data used for this study is not publicly available. Researchers interested in a collaboration are welcome to contact the senior authors (Prof. Jeroen Van Schependom and Prof. Guy Nagels).

## Code Availability

The analyses were performed exclusively in Python using the osl-ephys (version 1.0.0) and osl-dynamics (version 1.4.5) packages. Both Python packages are publicly accessible on the OHBA Analysis Group GitHub: https://github.com/ohba-analysis. The contents of the performed analysis are based on the methodology reported by Quinn et al.^15^ and Rossi et al.^22^, together with example codes for the data preprocessing (https://github.com/OHBA-analysis/osl-ephys/tree/main/examples) and the TDE-HMM analysis (https://github.com/OHBA-analysis/osl-dynamics/tree/main/examples). Analysis scripts written in Python (Jupyter Notebooks) are available upon request from the corresponding author.

## Acknowledgments

The authors would like to thank all participants, both healthy controls and people with multiple sclerosis, for their time, enthusiasm, and dedication to this research. The MEG data collection was enabled by a grant from the Belgian Charcot Foundation and an unrestricted research grant by Genzyme-Sanofi awarded to G.N., who is a senior clinical research fellow of the Fonds Wetenschappelijk Onderzoek (FWO) Flanders (1805620N). O.B. is funded by FWO, under grant number 1109225N.

