## Supplementary Information for "Weakened prefrontal activation dynamics associated with slowed information processing speed in multiple sclerosis"

#### 1. Bad channel and segment annotation

The OSL Python preprocessing pipeline includes an automated bad channel and segment identification step, which was not implemented by default in the legacy MATLAB pipeline[1]. For the dataset used in this work, the average proportion of identified bad channels across all subjects is 2.29 ± 0.02 %, and 1.17 ± 0.02 % for the magnetometer and gradiometer channels, respectively. Table S1 reports these proportions for different subgroups. There are no significant differences between healthy controls (HCs) and people with multiple sclerosis (PwMS), or between the two scanner types used for MEG data collection (original scanner 1 – 33 subjects, upgraded scanner 2 – 77 subjects).

| Type | HCs (%) | PwMS (%) | p-value | Scanner 1 (%) | Scanner 2 (%) | p-value |
|---|---|---|---|---|---|---|
| **Bad channel (mag)** | 2.28 ± 0.02 | 2.30 ± 0.02 | 0.705 | 1.87 ± 0.02 | 2.47 ± 0.02 | 0.187 |
| **Bad channel (grad)** | 1.00 ± 0.02 | 1.25 ± 0.02 | 0.316 | 1.59 ± 0.02 | 0.99 ± 0.02 | 0.251 |
| **Bad segment rejected** | 9.65 ± 0.07 | 10.23 ± 0.07 | 0.783 | 7.12 ± 0.07 | 11.28 ± 0.07 | **0.0006*** |

**Table S1 Bad channels and segments identified during MEG preprocessing**. The values are reported as average proportions (%) across all subjects belonging to a certain group (mean ± std). Bad channel proportions are expressed against the total number of channels (magnetometer = 102, gradiometer = 204), and the proportions of rejected epochs containing a bad segment are expressed against the total 128 trials. Group comparisons (HCs vs. PwMS, and Scanner 1 vs. Scanner 2) were conducted using the Mann-Whitney U-test. *$p < 0.05$.

When epoching the hidden state time course to obtain the state-wise temporal description, epochs containing a bad segment within their time window were rejected. Out of each participant's 128 trials, the average proportion of rejected epochs across all subjects is 10.23 ± 0.07 %. From Table S1, this proportion significantly differs based on the scanner type used to acquire MEG data (Mann-Whitney U-test, p-value = 0.0006). To verify that the scanner upgrade does not confound state inference and the task-related state activation profiles, we performed two separate TDE-HMM inferences on the subject groups divided by type of MEG scanner (see supplementary materials, section 9).

#### 2. Task performance

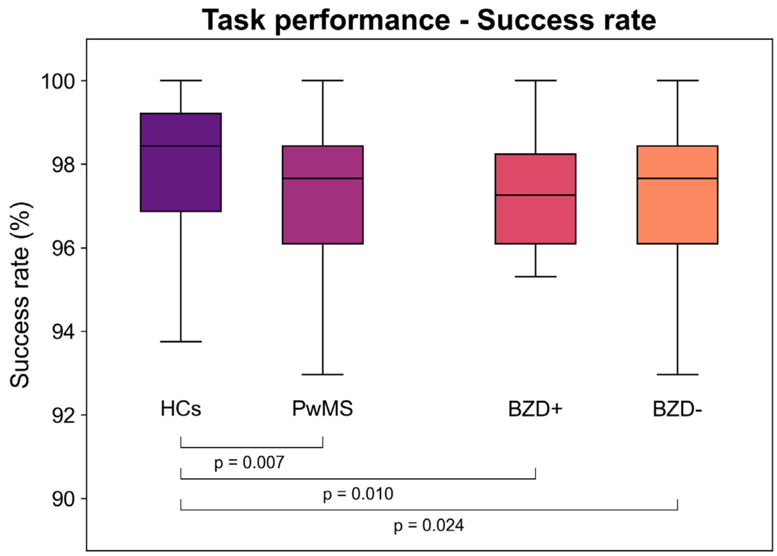


**Figure S1 Success rates distribution from the subjects' responses during the computerised SDMT.** Pressing the wrong button or not pressing the button at all during the maximum allowed response window (6 s) is considered a mistake. Group differences between the different subgroups (HCs, PwMS, BZD+ and BZD-) were performed using the Mann-Whitney U-test.

The distribution of success rates during the computerised SDMT (i.e. the accuracy of the given response) for different subgroups is shown in Figure S1. HCs have significantly higher success rates compared to PwMS (Mann-Whitney U-test, p-value = 0.00696). Overall, nearly all subjects only make a few or no mistakes throughout the total 128 trials.

Table S2 shows the mean reaction times (RTs) per paradigm condition (correct/incorrect). When subjects are presented with an incorrect stimulus (mean RT = 1.814 s), their average RT is slower compared to when correct stimuli are presented (mean RT = 1.668 s). Group comparisons for the different paradigm conditions (correct/incorrect) and response types (ok/mistake) are reported in Table S2. The RT significantly differs between HCs and PwMS for both correct stimuli (Mann-Whitney U-test, p-value = $4.9 \cdot 10^{-51}$) and incorrect stimuli (Mann-Whitney U-test, p-value = $1.7 \cdot 10^{-56}$), specifically for correct subject responses (response type = ok). Although the RTs for mistake responses also lead to p-values below the significance threshold, these results were not used to draw any conclusions given the very small occurrence of mistake responses. Considering the BZD+ and BZD- groups, only the RT for correct stimuli significantly differ between those two groups (Mann-Whitney U-test, p-value = 0.003), again only considering the correct subject responses (response type = ok).

| Stimulus | Response | HCs | PwMS | p-value | BZD+ | BZD- | p-value |
|---|---|---|---|---|---|---|---|
| **Correct** | **ok** | 1.52 ± 0.46 | 1.72 ± 0.52 | **$4.9 \cdot 10^{-51}$*** | 1.77 ± 0.55 | 1.72 ± 0.52 | **0.003*** |
| | **mistake** | 1.61 ± 0.48 | 1.91 ± 0.57 | **0.002*** | 1.62 ± 0.51 | 1.92 ± 0.57 | **0.006*** |
| **Incorrect** | **ok** | 1.65 ± 0.55 | 1.89 ± 0.66 | **$1.7 \cdot 10^{-56}$*** | 1.92 ± 0.67 | 1.89 ± 0.66 | 0.24 |
| | **mistake** | 1.71 ± 0.60 | 1.99 ± 0.71 | **0.029*** | 1.85 ± 0.57 | 1.99 ± 0.71 | 0.47 |

**Table S2 Mean reaction times (RTs) from the subjects' responses during the computerised SDMT.** Values are reported for the two paradigm conditions (correct/incorrect) and separated based on whether the subject pressed the appropriate button (ok/mistake) given the presented stimuli. RTs are expressed in seconds (mean ± std). Group differences between the different subgroups (HCs vs. PwMS, and BZD+ vs. BZD-) were evaluated using the Mann-Whitney U-test. All p-values were corrected for multiple comparisons through FDR correction. *p < 0.05.

### 3. Hyperparameters

| HMM process | Hyperparameter | Value |
|---|---|---|
| TDE data preparation | embeddings | 15 |
| | PCs for PCA | 80 |
| Model inference | hidden states | 6 |
| | random initialisations | 3 |
| | sequence length | 2000 |
| | learning rate | 0.01 |
| | batch size | 32 |
| | epochs | 30 |

**Table S3 Hyperparameters used for the data preparation and TDE-HMM inference process.** The number of time-delay embeddings and number of principal components (PCs) for the PCA reduction step are used to prepare the parcelled MEG data before serving as input for model inference. The number of hidden states, number of random initialisations to provide initial values to the model parameters, sequence length expressed in samples, learning rate, batch size, and number of epochs are relevant for the TDE-HMM inference. We chose these hyperparameters according to the work and recommendations of Vidaurre et al.[2] For the dataset used in this work, the loss function converged after about 30 epochs.

### 4. Summary statistics

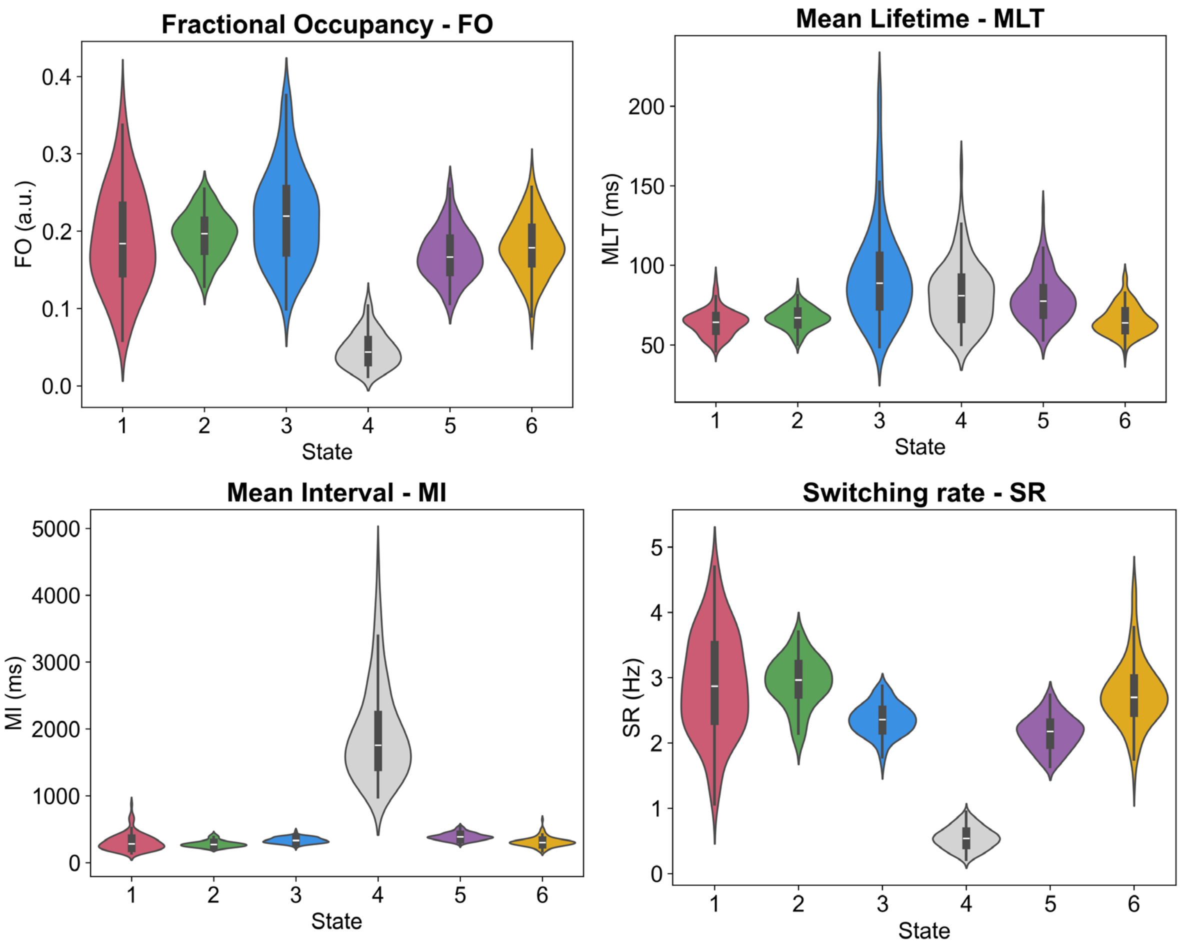


**Figure S2 Summary statistics for the extracted HMM states**. The states' temporal characteristics are summarised by four metrics: Fractional Occupancy (FO), Mean Lifetime (MLT), Mean Interval (MI), and Switching Rate (SR). Violin plots show the distribution of the state's temporal properties across all subjects.

Distributions of the four metrics (fractional occupancy, mean lifetime, mean interval, and switching rate) across all subjects per state are shown in Figure S2. The exact values of the population-level mean and standard deviation per metric and state are reported in Table S4. Considering the fractional occupancy (FO), the average FO per state is around 17%. State 3 occurs the most (22% of the time), while state 1, 2, 5 and 6 are activated for about 18% of the time. State 4 only activates for about 5% of the time. Interestingly, state 1 and state 3, the cognitive states, show the largest standard deviation in fractional occupancy. The mean lifetime (MLT) across states goes from a shorter 65 ms (state 1, 2, and 6) to a longer 95 ms (state 3, and 4). The mean interval (MI) for state 1, 2, 3, 5, and 6 are comparable and lie between 280 and 400 ms. State 4 has the longest average time duration between successive state visits, at 1938 ms. The switching rate (SR) lies between 2-3 Hz for state 1, 2, 3, 5, and 6. As state 4 has the highest MI, it makes sense that this state would also have the lowest SR of 0.54 Hz. Finally, possible significant differences in summary statistics between HCs and PwMS were investigated using a permutation test (N = 10 000) with the maximal statistic taken across states. No significant differences (p-value below the significance threshold of 5%) were observed.

| | FO | MLT (ms) | MI (ms) | SR (Hz) |
|---|---|---|---|---|
| State 1 | 0.19 ± 0.07 | 64.04 ± 8.14 | 310.37 ± 127.34 | 2.90 ± 0.77 |
| State 2 | 0.19 ± 0.03 | 66.77 ± 6.66 | 280.93 ± 48.92 | 2.93 ± 0.38 |
| State 3 | 0.22 ± 0.06 | 94.34 ± 30.69 | 334.93 ± 44.20 | 2.35 ± 0.24 |
| State 4 | 0.05 ± 0.02 | 94.34 ± 20.26 | 1938.31 ± 713.53 | 0.54 ± 0.16 |
| State 5 | 0.17 ± 0.03 | 79.39 ± 14.92 | 390.04 ± 55.7 | 2.16 ± 0.26 |
| State 6 | 0.18 ± 0.04 | 65.28 ± 9.09 | 309.30 ± 70.89 | 2.76 ± 0.51 |

**Table S4 Quantified summary statistics for the extracted HMM states.** The values are reported as averages of the states' properties across all subjects (mean ± std). Group comparisons (HCs vs. PwMS) were conducted using the permutation test with max t-statistic (N=1000), and resulted in no significant group differences.

### 5. Additional GLM contrasts – effect of covariates

Figure S3 reports the effect of several categorical regressors on the event-related state activation profiles. The stimulus type does not appear to modulate the activation profiles of the states. Considering that subjects were asked to press a button during each trial and did not receive feedback on their performance, this lack of effect between the two paradigm conditions seems reasonable. Looking at the disease effect plot, we observe several significant group differences between HCs and PwMS. Specifically, certain time windows in state 1 and 5 show significantly reduced event-related state activation in PwMS compared to HCs, while state 3 exhibits significantly increased activation in PwMS throughout nearly the entire epoch window. These results were reported (see Figures 4 and 5) and further discussed in the main text. Regarding the effect of benzodiazepine uptake, we observed significantly increased activation in state 3 for the BZD+ PwMS group, only between 1200 and 1500 ms post-stimulus. Considering state 3's characteristic beta activity (Figure 3b), this supports the known effect of benzodiazepines in enhancing beta activity.[3–5] Rossi et al. reported a similar amplification of frontoparietal beta activity in BZD+ PwMS during the n-back task.[6] Lastly, females exhibit increased activation in state 3, whereas males show increased activation in state 5 and 6. However, all of these effects are limited to only a small range of time points.

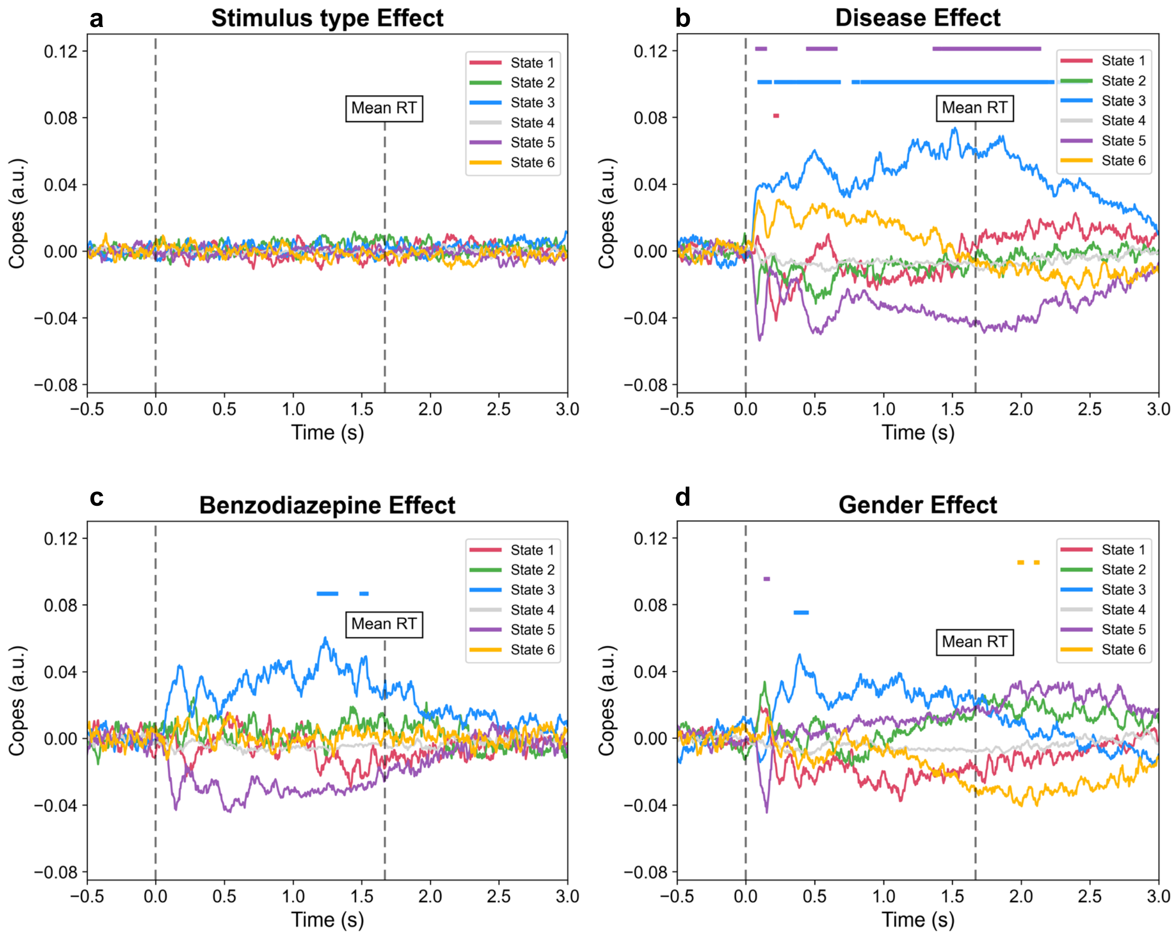


**Figure S3 Effect of categorical regressors on the state activation profiles.** In every subplot, each curve – associated to one state – represents the GLM contrast of parameter estimates (COPES), expressed as the difference between two conditions, considered within [-500, 3000] ms around the stimulus onset. The evaluated contrasts are: (a) stimulus type effect (incorrect – correct), (b) disease effect (PwMS – HCs), (c) benzodiazepine effect within the MS cohort (BZD+ – BZD-), and (d) gender effect (female – male). The horizontal lines at the top indicate time points of significant modulation of the state activation by the considered covariate, determined using a permutation test (N = 1000) with the maximum statistic (corrected for multiple comparisons across states and time points) at a significance level of 5%.

### 6. Spectral components

Figure S4 shows four spectral components obtained through data-driven spectral decomposition using non-negative matrix factorisation (NNMF). Spectral component 1 (blue) contains the delta (1-4 Hz) and theta (4-8 Hz) bands, forming the classical low-frequency bands. Spectral component 2 (yellow) represents the alpha band (8-13 Hz), and spectral component 3 (green) mainly captures the beta band (13-30 Hz). The fourth spectral component (purple) has been reported to capture high-frequency noise and is usually omitted during the states' description.[7]

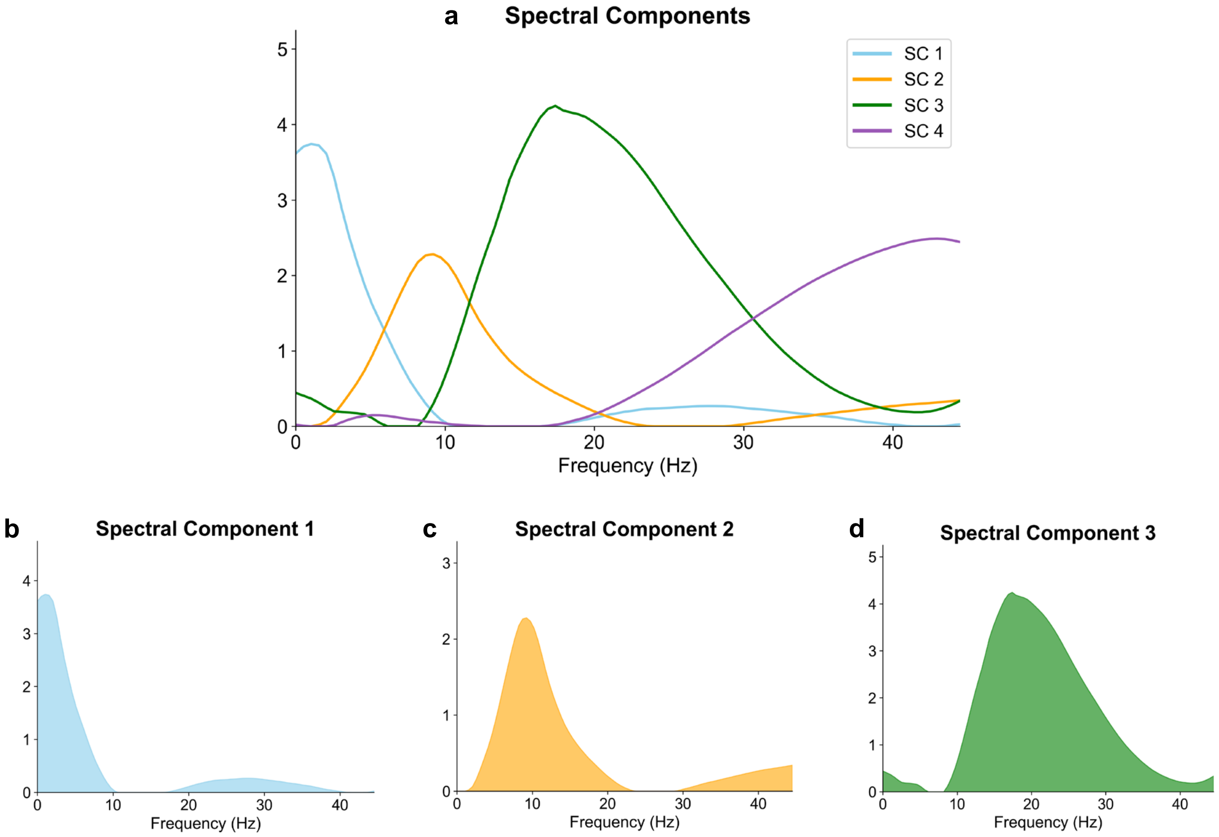


**Figure S4 Data-driven frequency bands (N=4) obtained by spectral decomposition using non-negative matrix factorisation (NNMF).** (a) All 4 extracted spectral components, (b) spectral component 1 – associated with the low-frequency bands, (c) spectral component 2 – represents the alpha band, (d) spectral component 3 – captures the beta band. The last spectral component (SC4) filters high-frequency noise and is omitted in the states' frequency content description.

#### 7. Complete spatio-spectral state descriptions

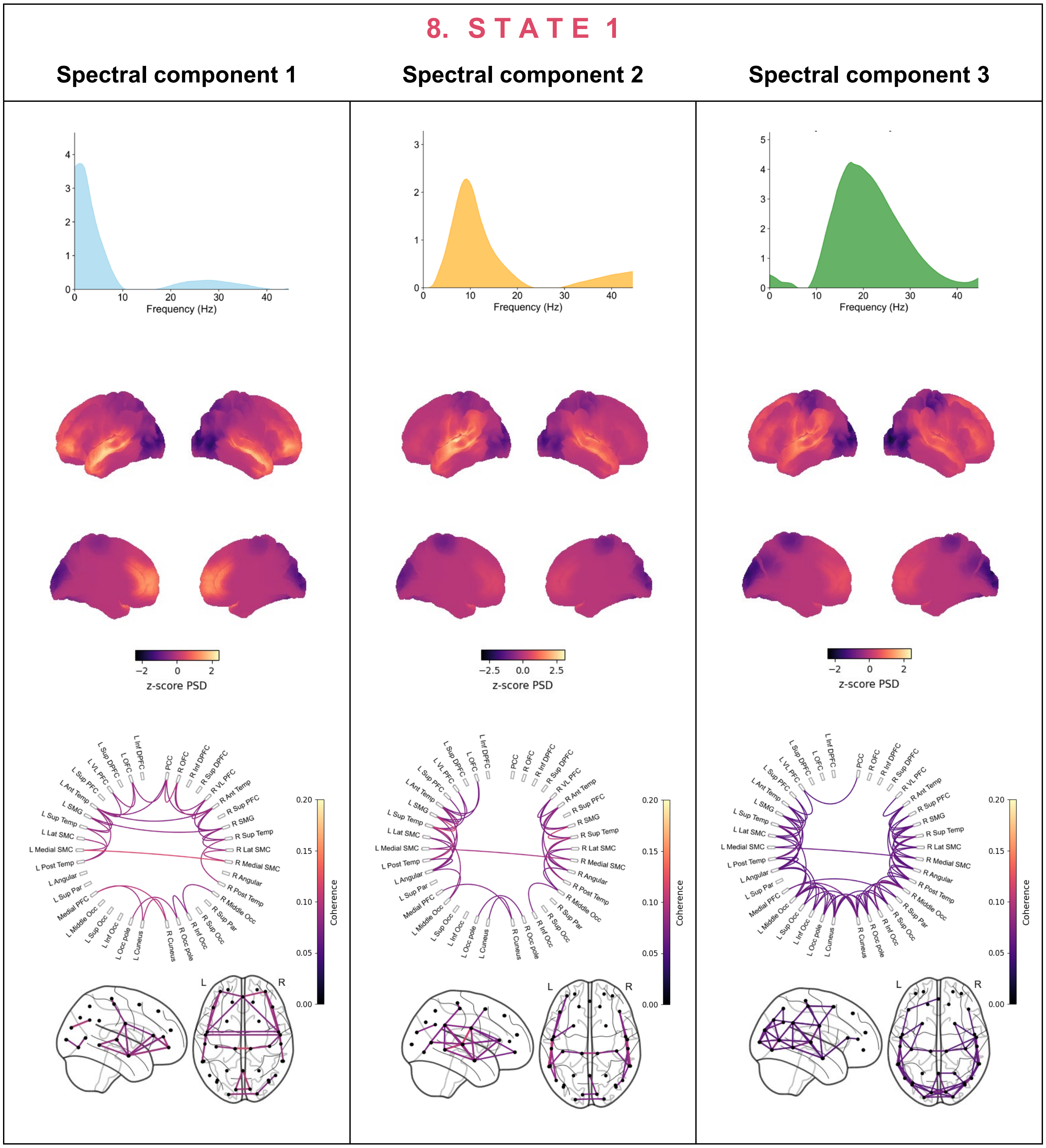


**Figure S5 Complete spectral description of state 1.** Every column provides the spatial and spectral state description weighted by one of the three spectral components. The first row shows the considered spectral component, while the second row contains the associated spatial power map, where the PSD is standardised (z-score) across parcels. The last row displays the coherence networks thresholded using a GMM, shown on a circular plot and a brain mask.

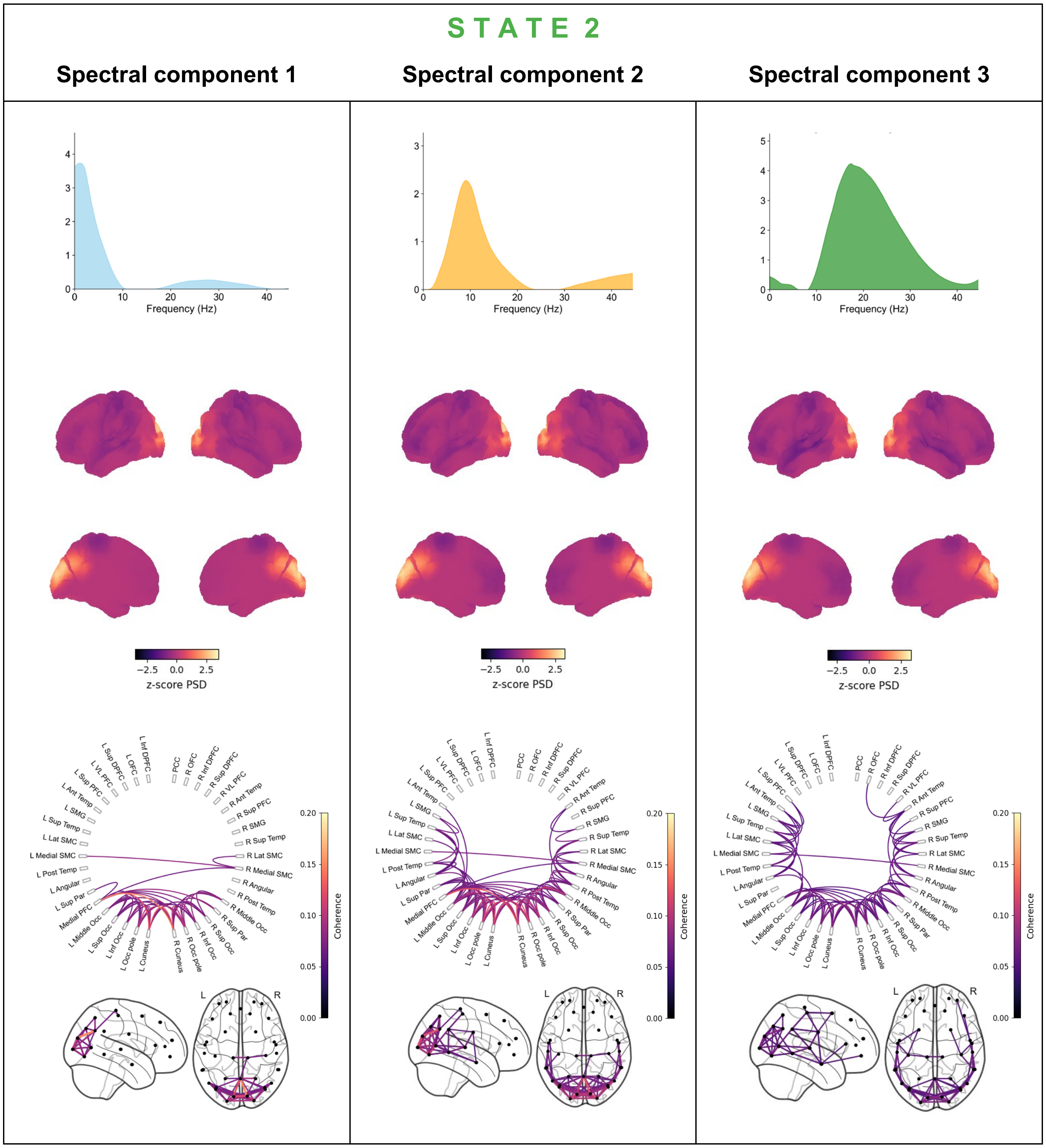


**Figure S6 Complete spectral description of state 2.** Every column provides the spatial and spectral state description weighted by one of the three spectral components. The first row shows the considered spectral component, while the second row contains the associated spatial power map, where the PSD is standardised (z-score) across parcels. The last row displays the coherence networks thresholded using a GMM, shown on a circular plot and a brain mask.

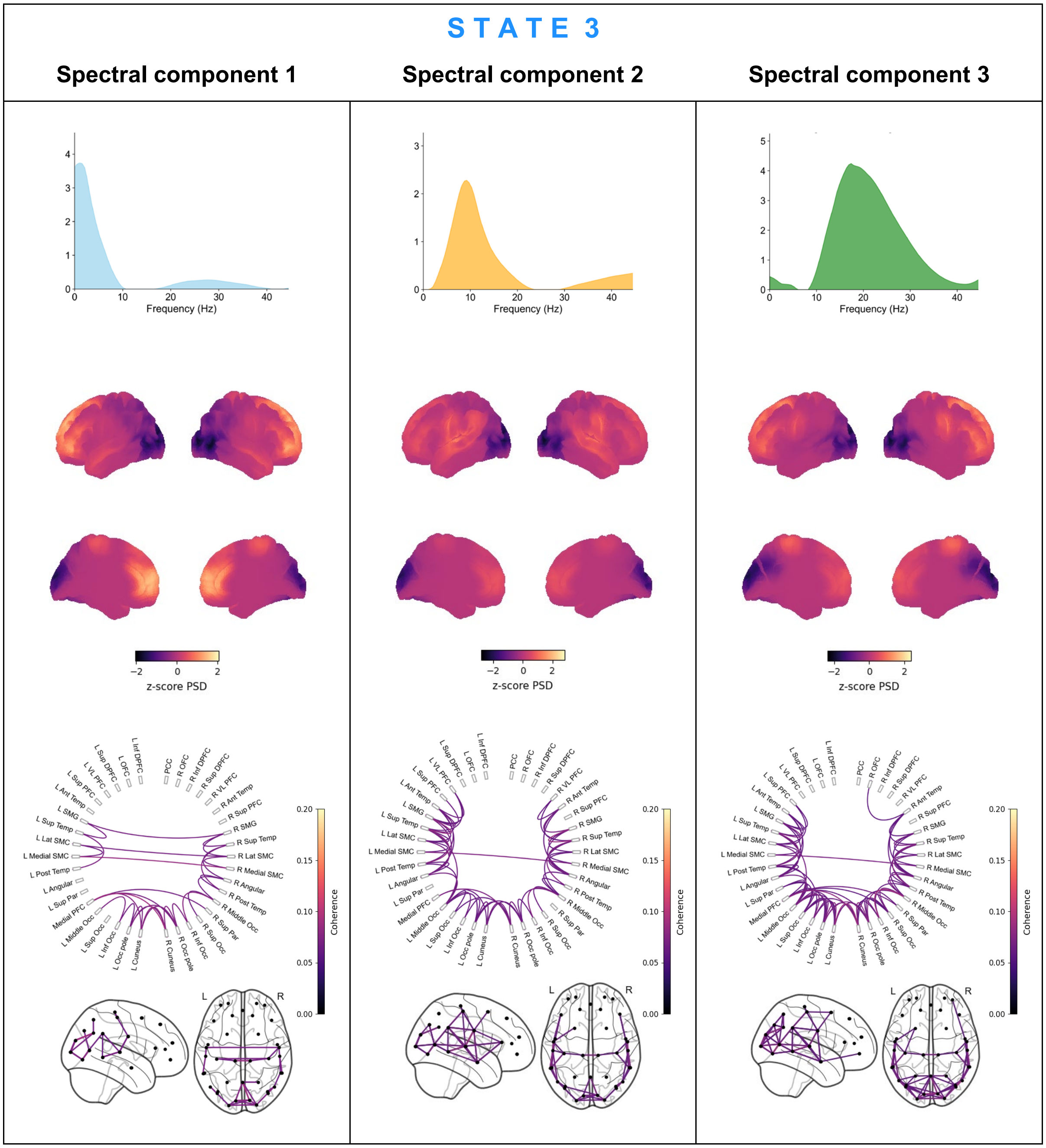


**Figure S7 Complete spectral description of state 3.** Every column provides the spatial and spectral state description weighted by one of the three spectral components. The first row shows the considered spectral component, while the second row contains the associated spatial power map, where the PSD is standardised (z-score) across parcels. The last row displays the coherence networks thresholded using a GMM, shown on a circular plot and a brain mask.

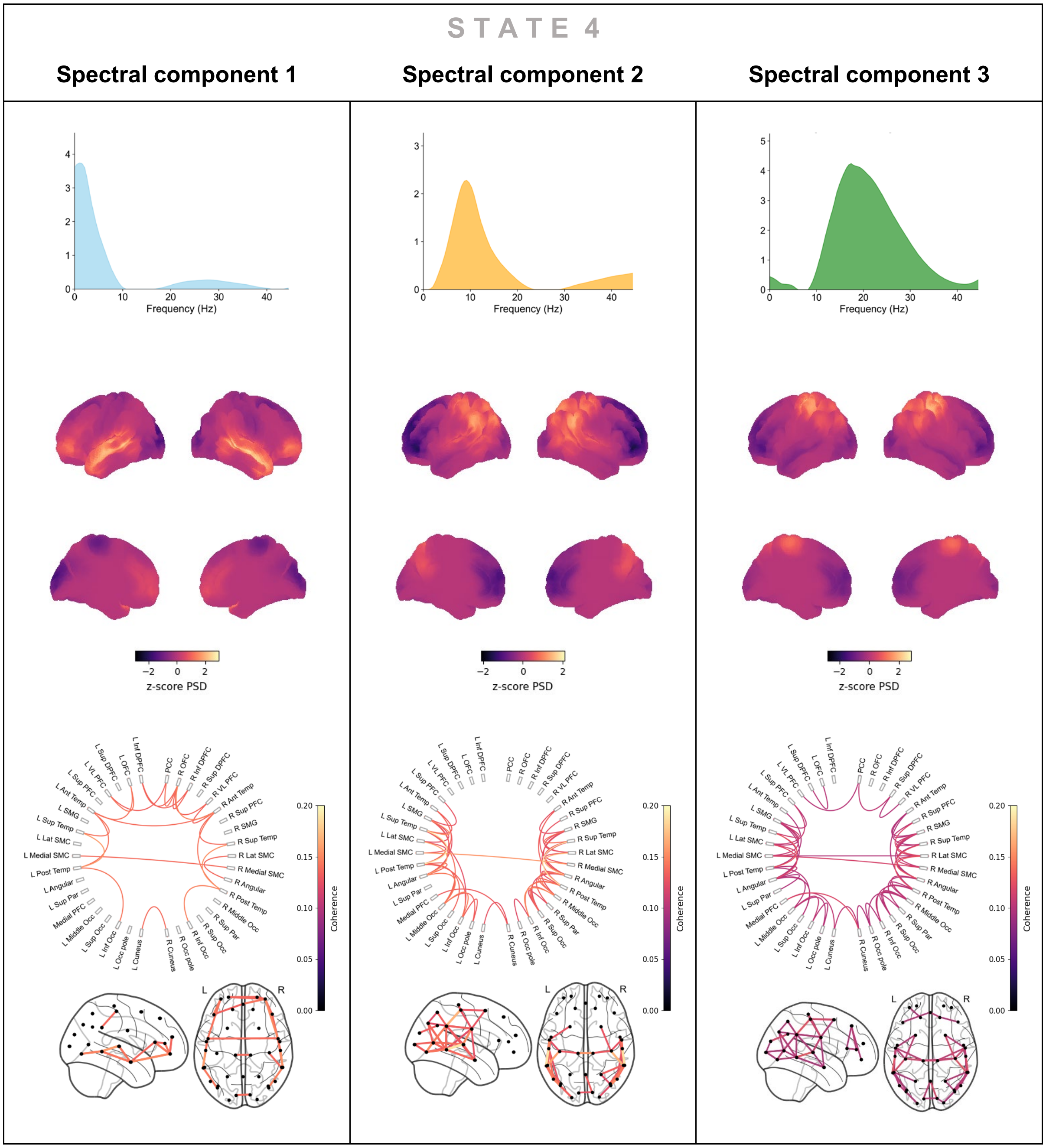


**Figure S8 Complete spectral description of state 4.** Every column provides the spatial and spectral state description weighted by one of the three spectral components. The first row shows the considered spectral component, while the second row contains the associated spatial power map, where the PSD is standardised (z-score) across parcels. The last row displays the coherence networks thresholded using a GMM, shown on a circular plot and a brain mask.

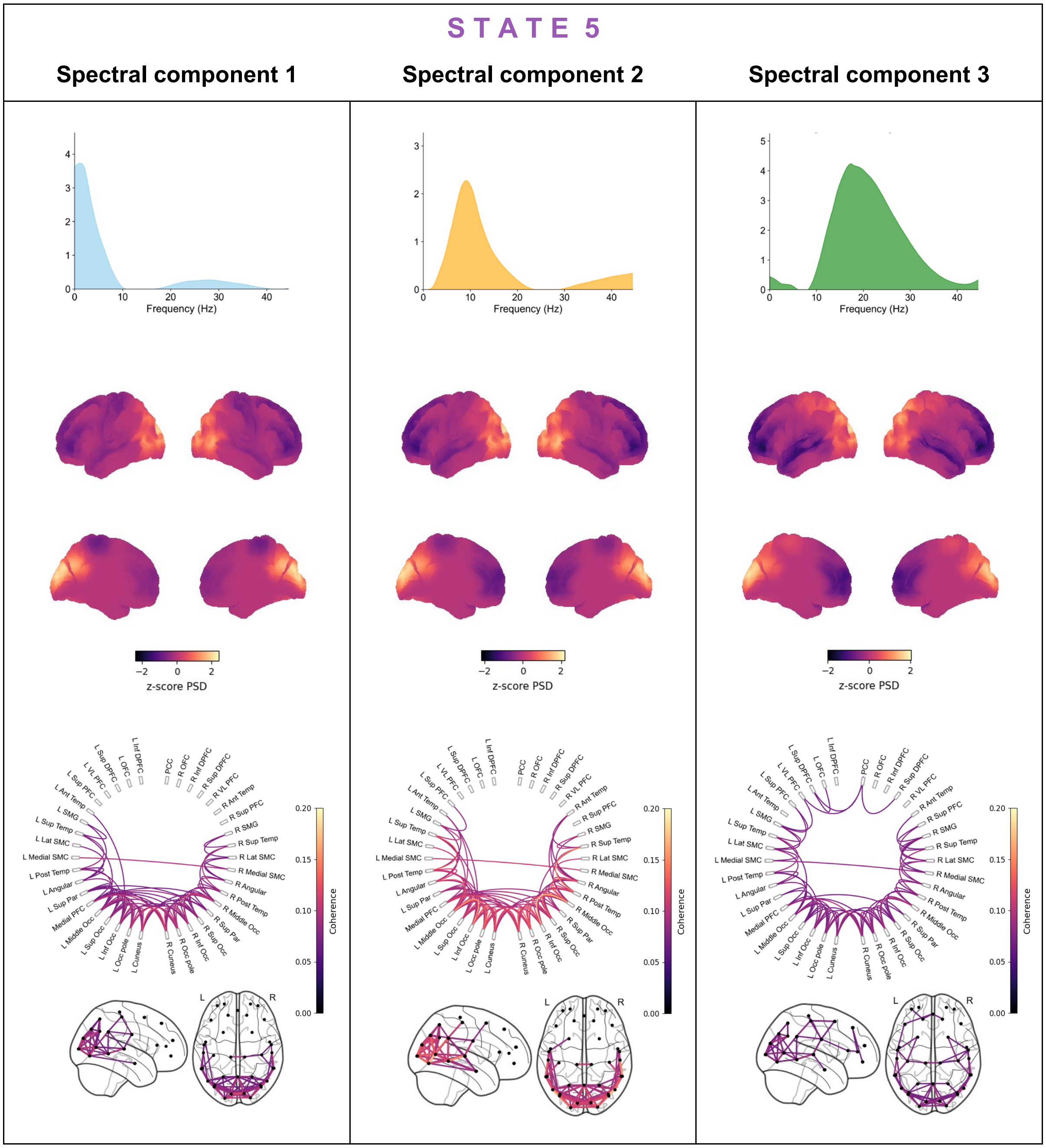


**Figure S9 Complete spectral description of state 5.** Every column provides the spatial and spectral state description weighted by one of the three spectral components. The first row shows the considered spectral component, while the second row contains the associated spatial power map, where the PSD is standardised (z-score) across parcels. The last row displays the coherence networks thresholded using a GMM, shown on a circular plot and a brain mask.

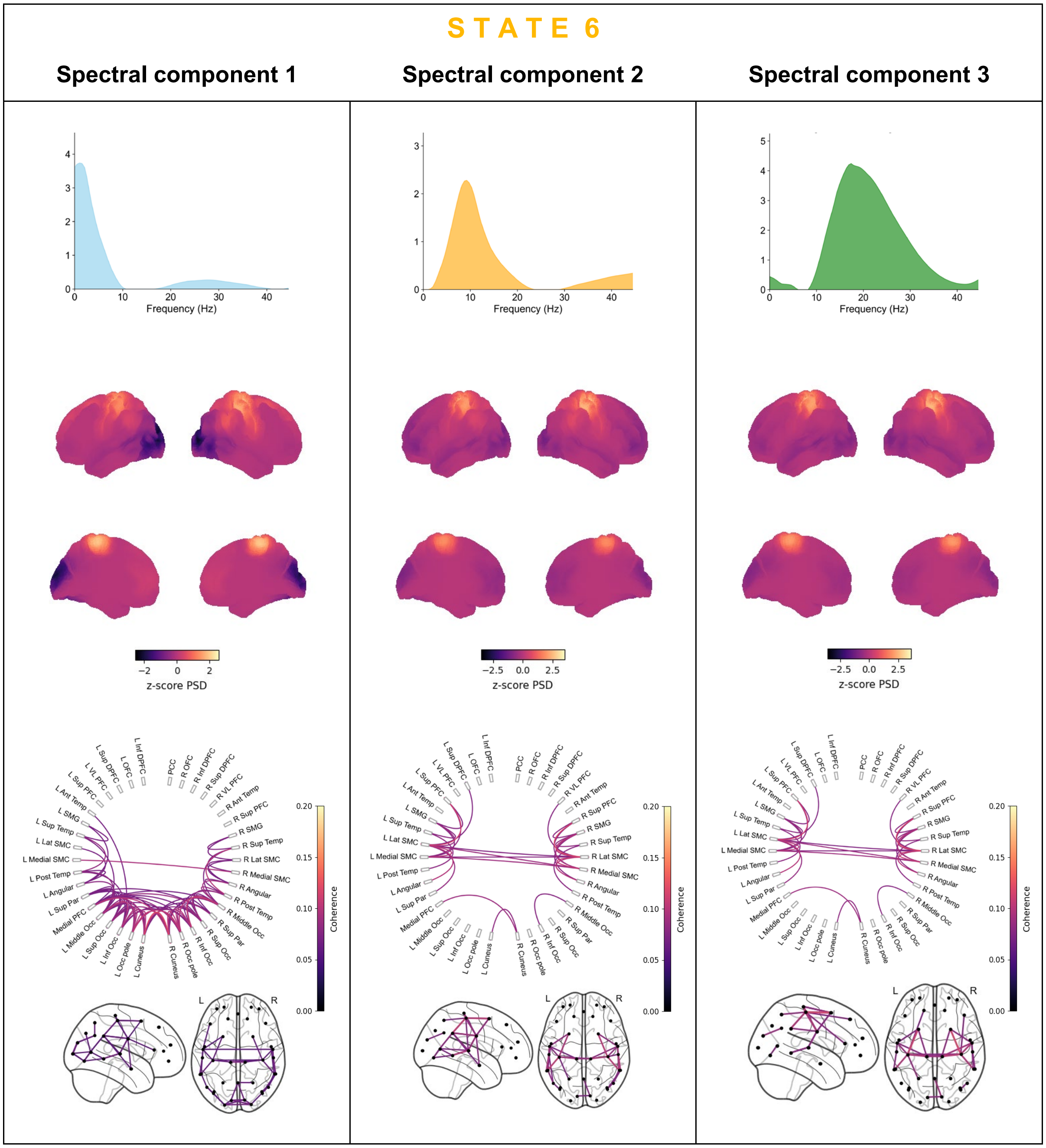


**Figure S10 Complete spectral description of state 6.** Every column provides the spatial and spectral state description weighted by one of the three spectral components. The first row shows the considered spectral component, while the second row contains the associated spatial power map, where the PSD is standardised (z-score) across parcels. The last row displays the coherence networks thresholded using a GMM, shown on a circular plot and a brain mask.

### 9. Additional Group comparisons – effect of Benzodiazepines and Gender

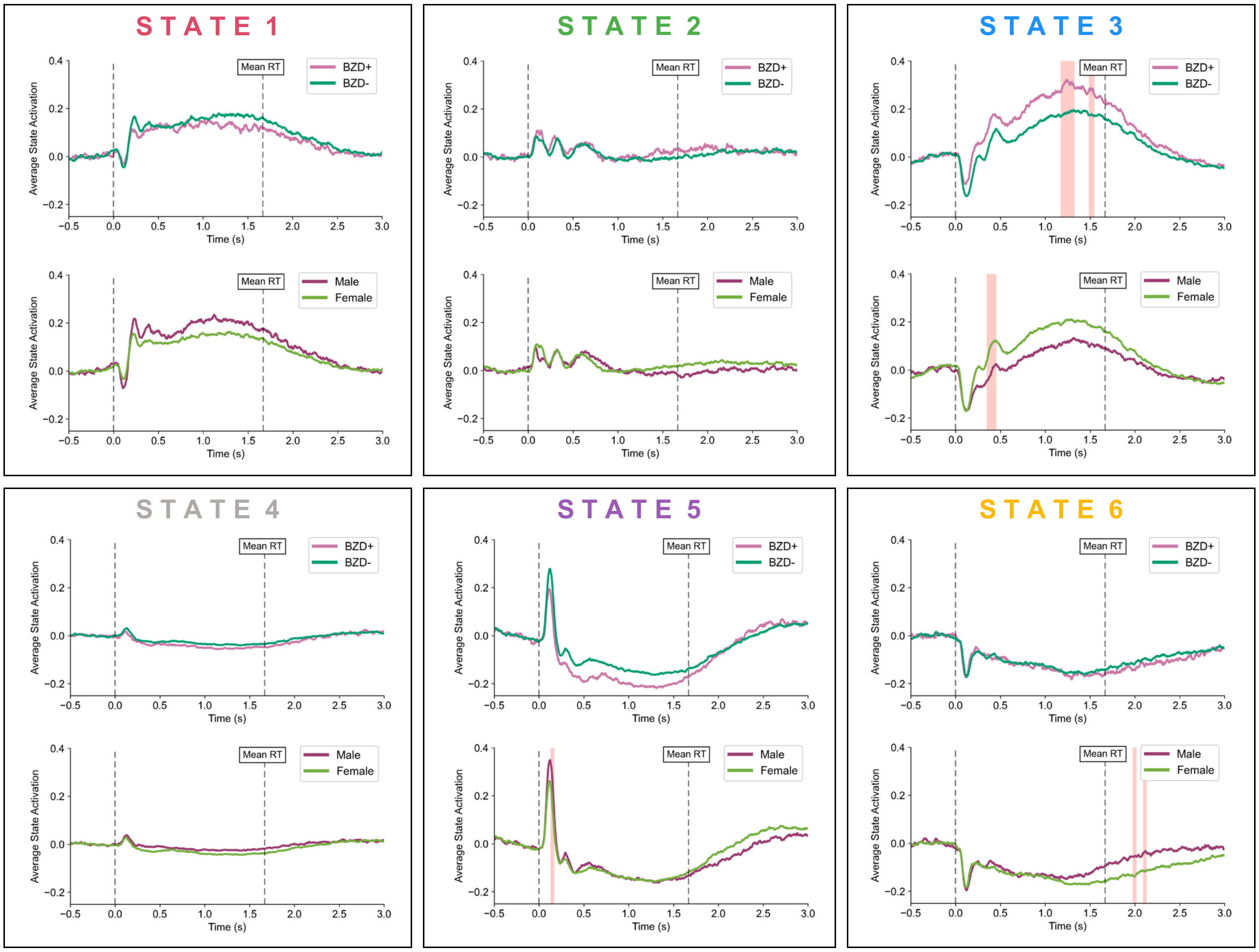


**Figure S11 Additional group comparisons for the effect of benzodiazepines and gender on the states' activation profile.** Each box is assigned to one of the states from the main analysis (6-state model) and displays two comparisons: benzodiazepine effect (BZD+ vs. BZD- PwMS) and gender effect (male vs. female), with solid lines representing the group-averaged epochs. Dotted vertical lines indicate the stimulus onset and mean reaction time, while pink shaded areas highlight time points where GLM contrasts reveal significant differences across groups (max t-stat permutation, $N = 1000$, $p < 0.05$).

### 10. Scanner type-specific TDE-HMM inferences

We ran two separate TDE-HMM inferences for subjects grouped by MEG scanner type (scanner 1 and scanner 2). When running several HMMs independently, the state labels and their associations with specific functional brain networks are assigned in an arbitrary order. By qualitatively comparing the state descriptions from both inferences (Figures S12 and S13), we successfully mapped all 6 states between the two inferences based on their highly similar temporal, spatial, and spectral characteristics. This mapping is reported in Table S5.

| Description | Scanner 1 | Scanner 2 |
|---|---|---|
| Occipital stimulus detection | State 2 | State 3 |
| Prefrontal | State 1 | State 2 |
| Occipital visual processing | State 4 | State 5 |
| Frontoparietal attention shift | State 5 | State 4 |
| Sensorimotor | State 3 | State 1 |
| Baseline (non-task-related) | State 6 | State 6 |

**Table S5 Mapping similar HMM states between two inferences.** The subjects whose data was used for each inference were grouped by MEG scanner type (scanner 1 and scanner 2). Both models were trained using the same hyperparameters as in the main analysis (see supplementary materials, section 3).

Regarding group differences in the temporal state activation profiles between HCs and PwMS, the same effects emerge as in the main analysis. These include, among others, a reduced early activation peak of the prefrontal network and an overall increase in frontoparietal network activation, both in PwMS. However, some of these effects remain non-significant across both inferences, likely due to limited statistical power, particularly in the scanner 1 group (33 subjects). Moreover, a sufficiently large sample size is crucial for improving the stability of group-level HMM inference.

### State inference for Scanner 1 group

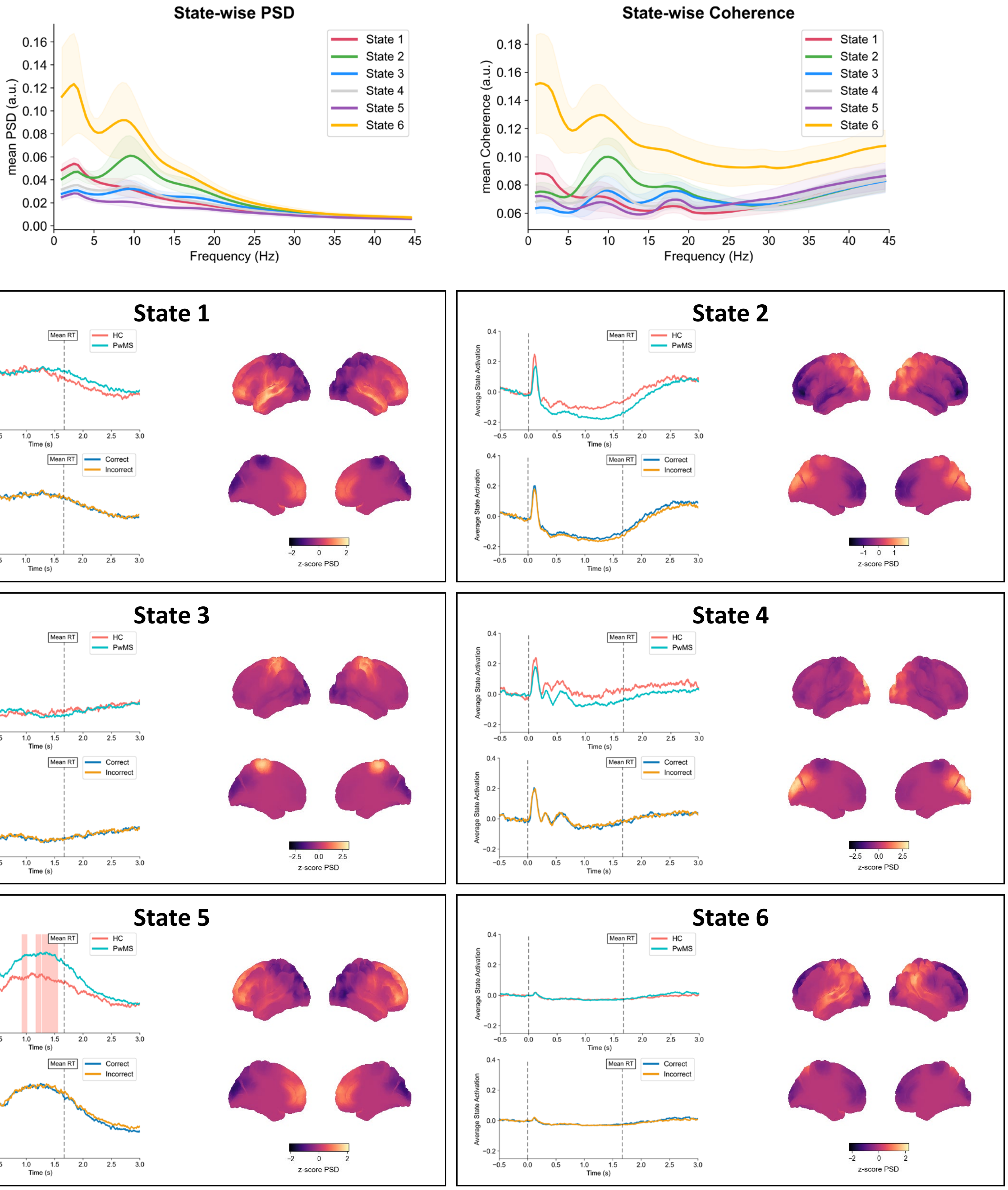


**Figure S12 State descriptions of the model inference in scanner 1 group.** At the top, we plot the mean power spectral density (PSD) and mean coherence for all states as a function of frequency (broadband), obtained by averaging across subjects, and all regions (PSD) or pairs of regions (coherence). Shaded areas at each frequency bin correspond to the standard deviation across subjects. Then, each box displays the temporal and spatial description of a state. Regarding the temporal activation profile, we display two comparisons: disease group (HCs vs. PwMS) and paradigm condition (correct vs. incorrect), with solid lines representing the group-averaged epochs. Pink shaded areas highlight time points where GLM contrasts reveal significant differences across groups or task conditions (max t-stat permutation, N = 1000, $p < 0.05$). The group-level average power is displayed on a spatial map, computed across the broadband frequency range [1-45] Hz, and standardised (z-score) across parcels.

### State inference for Scanner 2 group

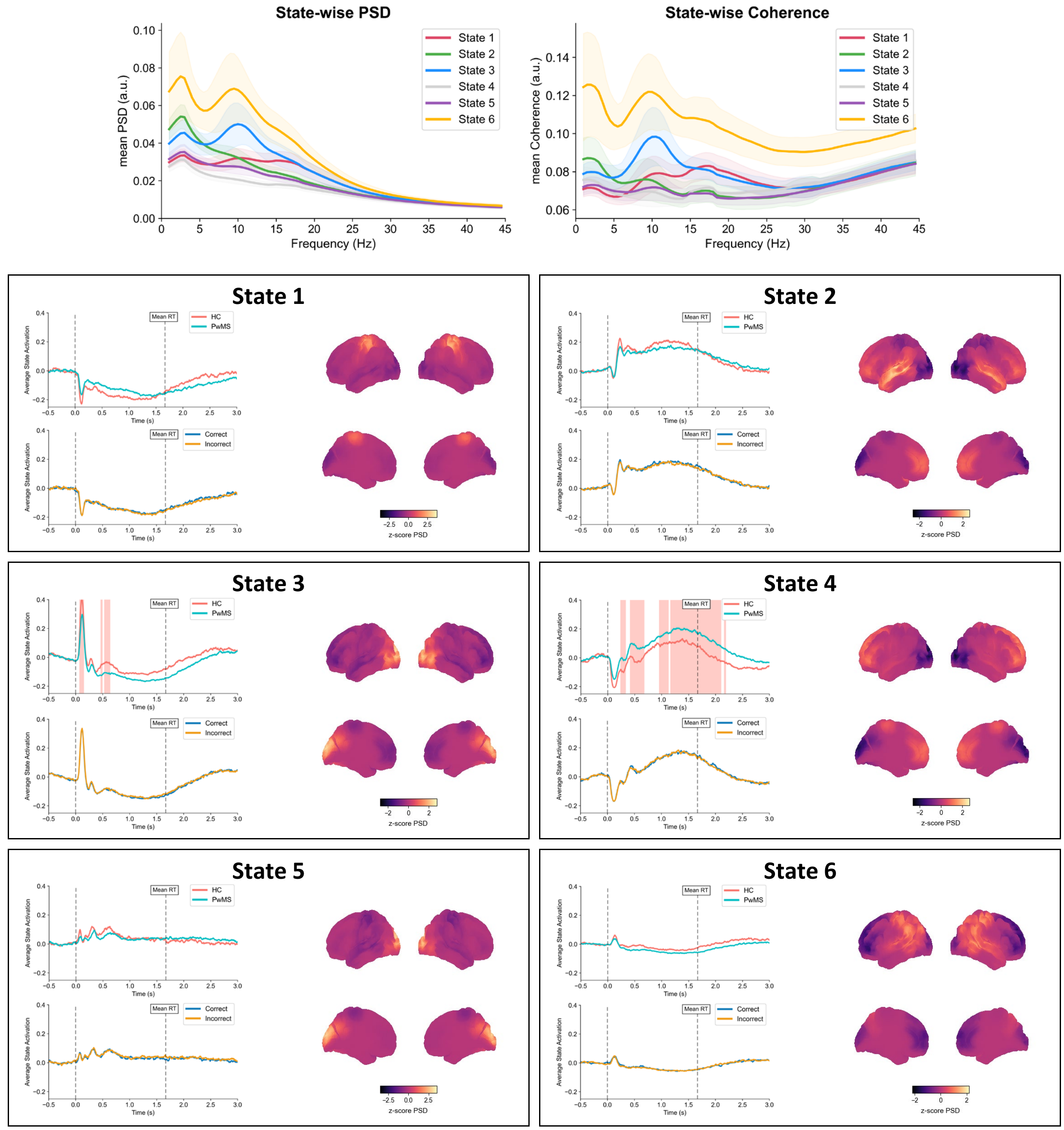


**Figure S13 State descriptions of the model inference in scanner 2 group.** At the top, we plot the mean power spectral density (PSD) and mean coherence for all states as a function of frequency (broadband), obtained by averaging across subjects, and all regions (PSD) or pairs of regions (coherence). Shaded areas at each frequency bin correspond to the standard deviation across subjects. Then, each box displays the temporal and spatial description of a state. Regarding the temporal activation profile, we display two comparisons: disease group (HCs vs. PwMS) and paradigm condition (correct vs. incorrect), with solid lines representing the group-averaged epochs. Pink shaded areas highlight time points where GLM contrasts reveal significant differences across groups or task conditions (max t-stat permutation, N = 1000, $p < 0.05$). The group-level average power is displayed on a spatial map, computed across the broadband frequency range [1-45] Hz, and standardised (z-score) across parcels.
